# Therapeutic Stress-induced Activation of PGCC Life Cycle Drives the Resistance Acquisition and Structured Tissue Differentiation

**DOI:** 10.64898/2026.04.04.716460

**Authors:** Zhiqian Zhang, Xiaoran Li, Xinxin Tian, Limin Deng, Jin-Tang Dong, Jinsong Liu

## Abstract

To elucidate how cancer cells generate resistance and defined forms of tissue structure in response to therapeutic stress, we tracked the temporal dynamics of life cycle of polyploid giant cancer cells (PGCCs) induced by the mitotic destabilizer vincristine (VCR). Live-cell fluorescence imaging revealed that VCR activated a distinct endoreplication-based life cycle, which replaced canonical mitosis. PGCCs exhibited reduced proliferative activity, enhanced epithelial-mesenchymal transition (EMT), progressive acquisition of blastomere-like features, and broad multilineage differentiation potential. Both PGCC populations and single PGCC-derived progeny displayed time- and dose-dependent acquisition of malignant traits *in vitro* and tumorigenic capacity *in vivo.* PGCC-derived spheroids exhibited ability to differentiate into the cells of origin from three germ layers. Importantly, pre-budding PGCCs induced by higher VCR concentrations exhibited enhanced ability for glandular structure formation and tissue differentiation. Morphologically, the nuclei of PGCC-derived spheroids underwent growth in size, formation of luminal structure, and followed by maturation of lumen. Mechanistically, PGCCs entered a senescent state characterized by elevated senescence-associated secretory phenotype (SASP)-manifested by rich proinflammatory cytokines. Notably, silencing IL1β, IL6, or IL8, or pharmacological inhibition of their receptors, suppressed PGCC formation, budding, EMT, and blastomere-like reprogramming into structured tissue. Our studies provide novel mechanistic insights into early embryogenesis and tumorigenesis at the tissue structural developmental level.

## 1. Introduction

The deep association between tumors and developmental differentiation fundamentally reflects the capacity of malignant cells to progress through developmental mimicry. Solid tumors manifest as diverse morphologically defined tissue architectures, including glandular, papillary, cystic, and solid patterns. Many high-grade cancers exhibit an undifferentiated phenotype resembling blastomere-stage embryonic cells, characterized by a high nuclear-to-cytoplasmic (N/C) ratio and primitive tissue organization. In contrast, low-grade or benign tumors often morphologically resemble later stages of organ maturation, displaying more differentiated glandular structures and reduced N/C ratios. Although these morphological hallmarks have been recognized by pathologists for over a century, the mechanistic basis underlying tumor tissue architecture remains poorly understood ^1–5^.

Polyploid giant cancer cells (PGCCs) represent a distinct subpopulation of enlarged tumor cells characterized by polyploid genomes and remarkable cellular plasticity within cancer tissues. PGCCs are detected in approximately 37–50% of human cancers and strongly correlate with disease relapse and poor clinical outcomes ^6,7^. Notably, their prevalence increases markedly in post-treatment specimens ^8^. Once dismissed as biologically inert or terminally senescent byproducts, PGCCs are now recognized as dynamic, stress-responsive entities that contribute to tumor initiation, recurrence, metastasis, and therapy resistance ^1,7,9,10^. This paradigm shift has positioned PGCCs as a powerful model for understanding cancer cell evolution and stress-adaptive tissue remodeling ^3,11^.

Following chemotherapy-induced stress, PGCCs frequently adopt a senescence-like state characterized by Senescence-Associated β-galactosidase (SA-β-gal) activity and robust activation of the senescence-associated secretory phenotype (SASP) ^12–14^. Remarkably, a subset of these cells can subsequently escape senescence through depolyploidization, a process involving asymmetric nuclear budding or reductive division that generates smaller, genomically reconfigured progeny with enhanced malignant potential ^13,15^. This bidirectional transition between senescence and depolyploidization provides an effective survival strategy under therapeutic pressure, endowing tumors with resilience and evolutionary adaptability.

In addition, PGCCs commonly exhibit epithelial–mesenchymal transition (EMT) features that further enhance their plasticity and invasive capacity ^16–19^. Intriguingly, our prior work suggests that PGCCs activate developmental programs reminiscent of the blastomere stage—an early pre-gastrulation state preceding canonical EMT activation. PGCCs transiently express pluripotency-associated markers, including OCT4, SOX2, and NANOG, and demonstrate multilineage differentiation potential ^20–26^. These observations suggest that PGCCs may undergo a blastomere-like nuclear reprogramming process in response to stress, recapitulating pre-implantation embryonic dynamics normally suppressed in somatic tissues. Reactivation of this latent developmental program may underlie the emergence of cellular heterogeneity, metastatic competence, and therapy resistance—hallmarks of advanced malignancy ^8,10^.

The discovery of pre-embryonic plasticity in PGCCs raises several fundamental questions. Does the PGCC life cycle serve as a driving force for stress-induced tissue regeneration that gives rise to the diverse tumor architectures observed histologically? How does the intensity of chemotherapeutic stress influence the fate of reprogrammed cells? And what is the mechanistic relationship among senescence, EMT, and structured tissue regeneration?

In the present study, we address these questions by spatially and temporally tracking the PGCC life cycle using gradient concentrations of vincristine (VCR), a microtubule-destabilizing agent and potent inducer of PGCC formation. Our findings establish a unified conceptual framework linking therapeutic stress intensity to PGCC fate bifurcation and tissue differentiation through coordinated activation of the senescence-associated secretory phenotype and the EMT programs.

## 2 Results

### 2.1 Temporal view of the life cycle of polyploid giant cancer cells

Our previous studies and from others have shown that PGCCs are inducible in cancer cell lines by hypoxia mimetics (e.g., CoCl), microtubule stabilizers (e.g., paclitaxel), or high-dose DNA-damaging agents ^8,17,21,22,27,28^. Yet, effects of microtubule destabilizers on PGCC formation/phenotype remain unexplored. Here, we used VCR, a microtubule destabilizer, to induce polyploidization in ovarian cancer cells. Using HEY (histone 2B-GFP labeled) and SKOV3 cells, we first profiled the full life cycle of VCR dose-dependent PGCCs. Briefly, both cell lines were treated with VCR at concentrations of 0.5, 1, and 4 μM for 2 days, followed by recovery in drug-free culture medium to facilitate the formation of PGCCs. Flow cytometric analysis demonstrated that the proportion of PGCCs increased with escalating VCR concentrations in both HEY and SKOV3 cells (**Fig. 1A-B, Fig. S1A-B, and Fig. S2A-C**).

**Figure 1.**
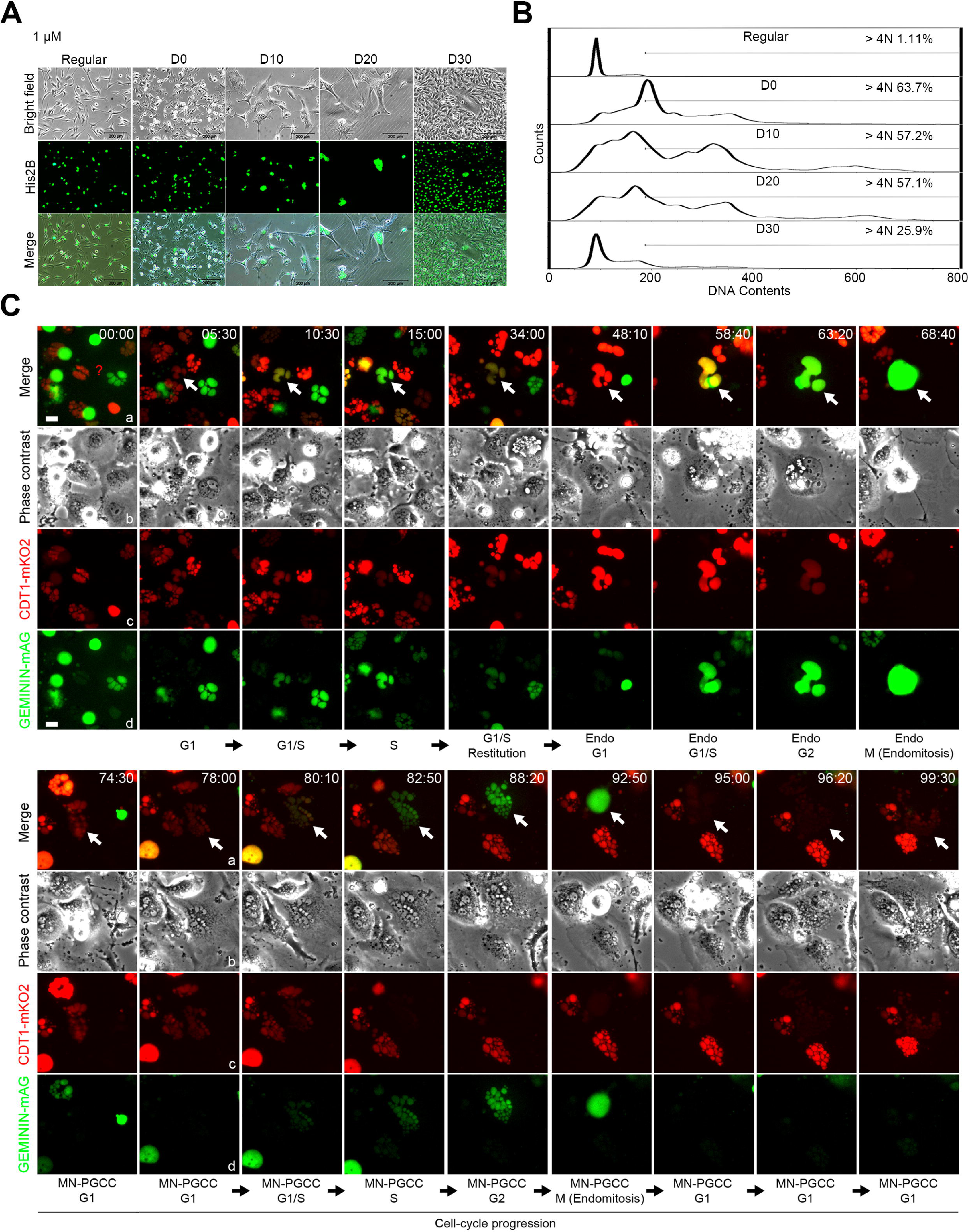
VCR induces the formation, endoreplication and budding processes of PGCCs in HEY cells. **(A and B)** The HEY cells were treated with 1 μM VCR for 2 days, then allowed to recover for PGCC formation and budding into daughter cells. A: Morphology of PGCCs at different time points. Upper lane: phase contrast images; middle lane: histone 2B-GFP labeled to show the nuclei; lower lane: merged images. B: Cell cycle distribution and percentage of cells with a DNA content greater than 4N in HEY cells treated with 1 μM VCR for 2 days and then allowed to recover in drug-free culture medium for the indicated days. Scale bars: 200 μm. **(C)** FUCCI-based live-cell imaging of VCR-induced PGCC formation. A stable HEY cell line expressing the FUCCI system was established to monitor cell cycle progression. Scale bars: 20 μm.

Meanwhile, HEY-derived PGCCs began to bud approximately 15-20 days post-recovery when treated with 0.5 μM VCR, whereas those exposed to 1 μM or 4 μM VCR initiated budding 20-25 days after recovery (**Fig. 1A-B and Fig. S1A-B**). SKOV3-derived PGCCs began to bud approximately 10-15 days post-recovery when treated with 0.5 μM VCR, whereas those exposed to 1 μM or 4 μM VCR initiated budding 15 –20 days after recovery (**Fig. S2A-C**). This concentration-dependent delay in budding onset suggests that higher VCR concentrations prolong the duration of homeostasis in PGCCs (**Fig. 1A-B, Fig. S1A-B, and Fig. S2A-C**).

To determine how VCR dosage affects PGCCs’ return to mitosis, we subcultured an equal number of PGCCs, generated from cancer cells treated with escalating VCR concentrations, into 12-well plates on day 10 (D10, defined as 10 days post-recovery after 2 days of VCR exposure). Daughter cell clones were stained with crystal violet 30 days post-subculture. Quantitative analysis revealed that the proportion of budding-capable PGCCs was approximately 3.1 ± 0.4% in HEY cells treated with 0.5 μM VCR. In HEY cells exposed to 1 μM and 4 μM VCR, this proportion decreased to 1.6 ± 0.3% and 1.2 ± 0.1%, respectively (**Fig. S1C-D**). A consistent trend was observed in SKOV3-derived PGCCs (**Fig. S2D**). Taken together, these results demonstrate that elevated VCR concentrations impair the ability of PGCCs to resume mitosis.

### 2.2 VCR induces the formation of PGCCs via endoreplication

To visualize nuclear dynamics during polyploidization, we established a HEY cell line stably expressing the FUCCI system (fluorescent ubiquitination-based cell cycle indicator) to monitor cell cycle progression and induce PGCC formation through VCR treatment. Live-cell imaging revealed distinct nuclear remodeling patterns under VCR exposure(**Fig. 1C and Supplemental video 1**). Following VCR treatment for 2 days and recovery in drug-free medium, cells progressed from G1 to S (**Fig. 1C**, 00:00-15:00), bypassing mitosis between 34:00 and 48:10 and initiating endoreplication. During this phase, progressive DNA accumulation drove nuclear enlargement (**Fig. 1C**, 48:10-63:20). The first endomitotic event at 68:40 produced ∼20 fragmented nuclei (**Fig. 1C**, white arrows), which exhibited synchronized cell cycle progression (**Fig. 1C**, 78:00, G1; 80:10, G1/S transition; 82:50, S/G2). A subsequent endomitosis, occurring between 88:20 and 95:00, generated ∼50 fragmented nuclear clusters (**Fig. 1C**).

We also microscopically tracked nuclear-size dynamics and cell cycle progression in individual PGCCs over 300 hours, starting on D8, to monitor nuclear changes during quiescence. While prolonged G1/S arrest was observed in most cells (**Fig. S3A**), progressive nuclear enlargement (**Fig. S3A and S3C**) indicated ongoing DNA replication. Notably, several PGCCs exhibited endoreplication cycles characterized by repeated G1/S-to-G2/M transitions and complete replication cycles (**Fig. S3B**), resulting in nuclear expansion (**Fig. S3B and S3D**). Importantly, when this PGCC started to bud, it entered the next endoreplication cycle, with nuclear size continuing to increase, albeit at a slower rate, as a portion of the DNA content had to be partitioned to the daughter cells (**Fig. S3B and S3D**).

To explore replication machinery dynamics in quiescent PGCCs, we generated HEY stable lines expressing H2B-mCherry/α-tubulin-GFP. Normal cells underwent canonical mitosis: nuclear enlargement (DNA synthesis), envelope breakdown, spindle assembly, metaphase plate formation, and cytokinesis, with daughter cells matching parental nuclear size (**Supplemental video 2 and Fig. S4, left panel**). In PGCCs (**Supplemental video 3 and Fig. S4, right panel**), multinucleated PGCCs (MuPGCCs) had heterogeneous nuclei. Entering prophase, envelopes fragmented rapidly and condensed chromatin formed discrete chromosomes. VCR-blocked microtubules caused diffuse α-tubulin-GFP signals (*vs*. bipolar spindles in controls), alongside “multi-pocket” morphology (from detached microtubules) and occasional chromatin bridges. Post-telophase, diminished tubulin signals preceded giant nucleus recovery via restitution endomitosis.

### 2.3 PGCCs and their daughter cells progressively develop malignant phenotypes

We systematically characterized the phenotypic and molecular characteristics (proliferation, migration, EMT and stemness) of PGCCs and their daughter cells. To characterize the phenotype of progeny cells derived from different concentrations of VCR-induced PGCCs, we compared the morphology of these cells with parental HEY cells. Morphological analysis revealed that the majority of daughter cells derived from PGCCs induced by 1 μM or 4 μM VCR exhibited a round and flat morphology at the initial stage of their life cycle (**Fig. 2A**). Prolonged passaging led to spindle-like morphological changes resembling regular HEY cells (**Fig. 2A**). Both MTS and SRB assays demonstrated that daughter cells derived from PGCCs induced by 1 μM or 4 μM VCR exhibited significantly slower growth rates when passaged for three generations compared to regular cancer cells (**Fig. 2B and S5A**). Notably, progeny derived from high-concentration (4 μM) VCR-induced PGCCs showed markedly weaker proliferative capacity than those generated under low-concentration (0.5 and 1 μM) conditions (**Fig. 2B and S5A**).

**Figure 2.**
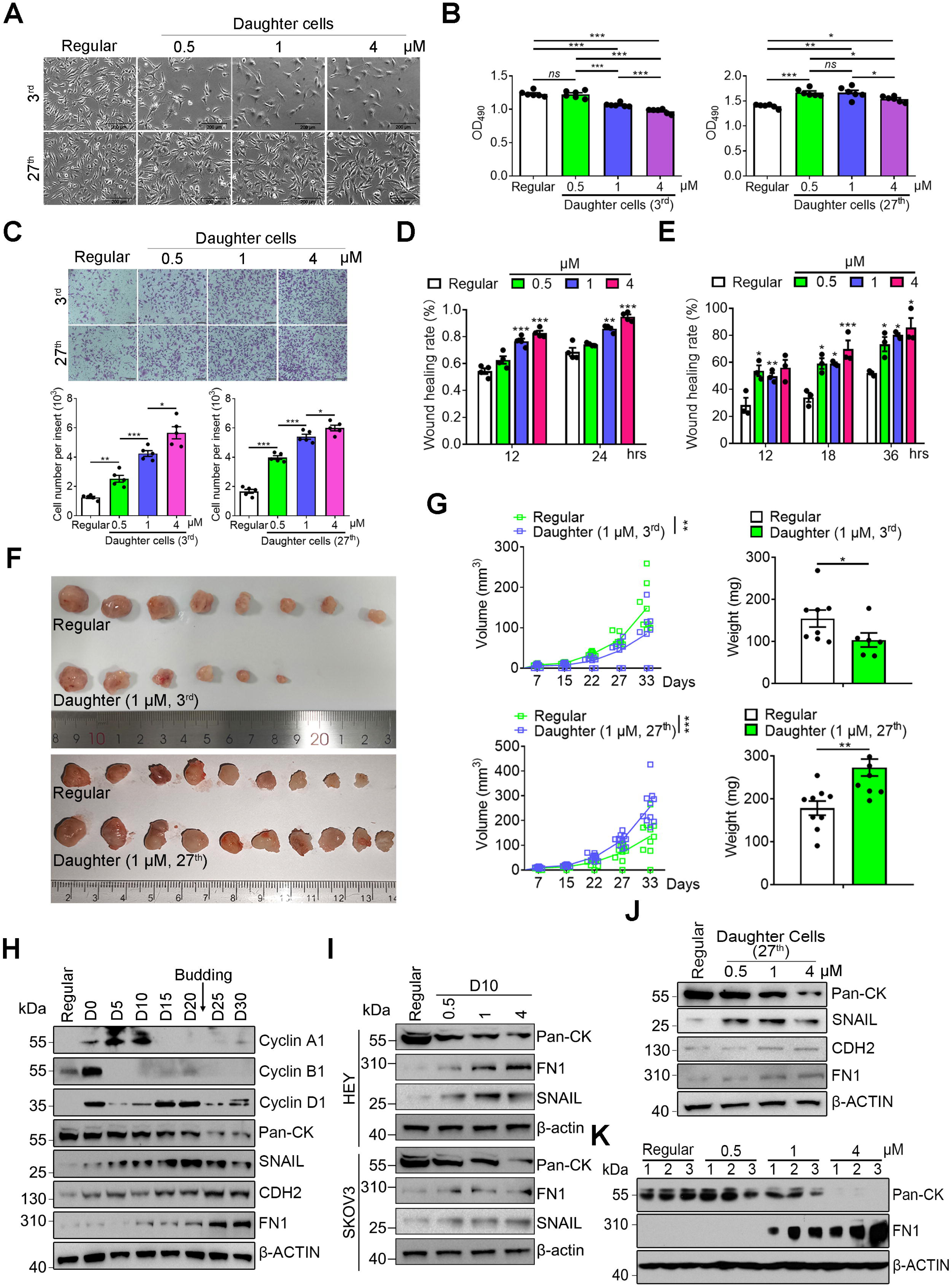
PGCCs and their progeny increasingly exhibit aggressive phenotypic traits over time. **(A)** Phase-contrast microscopy images of daughter cells derived from HEY cells induced by vincristine (indicated concentrations) at the 3^rd^ and 27^th^ passages, illustrating morphological alterations. Scale bars: 200 μm. **(B)** MTS experiments quantifying proliferation rates of 3^rd^- (left) and 27^th^- (right) passage daughter cells relative to regular HEY cells. Data are presented as mean ± SEM. *ns*, *p* > 0.05, * *p* < 0.05, ** *p* < 0.01, *** *p* < 0.001. **(C)** Transwell migration assay assessing the migratory capacity of 3^rd^- and 27^th^-passage daughter cells versus regular HEY cells. Upper layer: Representative images of migrated cells. Lower layer: Quantification: average cell counts per insert. Data are presented as mean ± SEM. *** *p* < 0.001. Scale bar: 200 μm. **(D)** Scratch wound healing assay demonstrating the migratory behavior of 27^th^-generation daughter cells derived from HEY cells induced by VCR at varying concentrations. Quantitative analysis of wound closure rates. Data are presented as mean ± SEM. \*\**p* < 0.01, *** *p* < 0.001. **(E)** Quantification of wound closure area in wound healing assay to evaluate the migratory capacity of progeny clones derived from different VCR-induced single PGCCs. Data are presented as mean ± SEM. * *p* < 0.05, ** *p* < 0.01, *** *p* < 0.001. **(F)** Xenograft assays were performed to evaluate the in vivo growth capacity of daughter cells derived from VCR-induced PGCCs (3^rd^ and 27^th^ generations) compared to regular HEY cells. **(G)** Growth curves (left) and tumor weights (right) of xenografts derived from 3^rd^ (upper) or 27^th^ (lower) generation daughter cells and regular cells are shown. Statistical analyses were performed using two-way ANOVA. Data presented as mean ± SEM. ** *p* < 0.01. *** *p* < 0.001. **(H)** Western blot analysis of Cell cycle-related proteins Cyclin A1, B1 and D1 and EMT-associated markers Pan-CK, SNAIL, CDH2, FN1 in HEY-derived PGCCs (1 μM VCR-induced) at sequential time points post-induction: D0, D5, D10, D15, D20, D25, and D30. **(I)** The levels of Pan-CK, FN1, and SNAIL in PGCCs at D10 induced by different concentrations of VCR and regular cancer cells were detected by Western blot analysis. Upper layer: HEY cells; Lower layer: SKOV3 cells. **(J)** Western blot analysis of EMT-associated markers Pan-CK, SNAIL, CDH2, and FN1 in 27^th^-passage daughter cells derived from different concentrations of VCR-induced PGCCs in HEY cells. **(K)** Western blot analysis of EMT markers Pan-CK and FN1 in daughter cell clones derived from different VCR-induced single PGCCs.

These data suggest that most daughter cells newly budded from PGCCs exhibit significantly reduced proliferative capacities. However, at the 27^th^ generation, daughter cells derived from PGCCs induced by low-concentration VCR (0.5 and 1 μM) exhibited significantly faster growth than those from regular cancer cells and the 4 μM cohort (**Fig. 2B and S5A**). This suggests that new-born daughter cells existed in a heterogeneous state with divergent proliferative capacities. Through continuous passaging, high-proliferative subpopulations gained dominance and ultimately hyperproliferative daughter cells became the dominant population that surpassed those of parental cancer cells. Also, we tested differences in drug resistance between the daughter cells and regular HEY cells. We found the daughter cells were more resistant to VCR than the regular HEY cells (**Fig. S5B**). Transwell migration assays further demonstrated that daughter cells derived from both the 3^rd^ and 27^th^ generations exhibited significantly enhanced migratory capacity compared to regular HEY cells (**Fig. 2C**). Results of the wound healing experiments also demonstrated that all daughter cells (27^th^ generation) filled the gap faster than the regular HEY cells (**Fig. S5C and Fig. 2D**). In addition, the daughter cells of PGCCs derived from higher VCR concentrations exhibited a higher gap-closing capability (**Fig. S5C and Fig. 2D**).

To exclude the possibility that the daughter cells originated from the re-proliferation of residual regular cancer cells during the formation of PGCCs, single PGCC-derived daughter cell clones were isolated and analyzed as shown in **Fig. S6A-B**. To determine the phenotype of these single-PGCC-derived daughter clones relative to the regular HEY cells, we randomly selected three clones from each VCR concentration group and performed wound-healing assays. Daughter clones demonstrated accelerated gap closure compared to the regular cells (*p* < 0.01), with the migration capacity exhibiting a concentration-dependent enhancement: clones derived from higher VCR concentrations displayed significantly greater migratory ability than those from lower concentrations (**Fig. S6C and Fig. 2E**).

To elucidate the growth capacity of PGCC-derived daughter cells *in vivo*, we established orthotopic xenograft models by subcutaneously implanting 5 × 10^6^ daughter cells budded from 1 μM VCR-induced PGCCs (3^rd^ and 27^th^ generations). alongside regular HEY cells into BALB/c nude mice. Mice inoculated with PGCC-derived daughter cells (3rd generation) displayed slower tumor growth kinetics **(Fig. 2F-G**, *p* < 0.01) and developed significantly lighter tumors compared to controls (**Fig. 2F-G**, *p* < 0.05). In contrast, mice inoculated with PGCC-derived daughter cells (^27^th generation) exhibited accelerated tumor growth kinetics (**Fig. 2F-G**, *p* < 0.001) and formed heavier tumors than control cohorts (**Fig. 2F-G**, *p* < 0.01). These results aligned with the *in vitro* findings (**Fig. 2B and S5A**), suggesting that continued in vitro passage of PGCCs-derived daughter cells allows selection of aggressive clones.

Finally, we analyzed the changes in cell cycle-related and EMT-associated proteins across phases of PGCCs’ life cycle. As shown in **Fig. 2H**, expression of cyclin A1, an S-to-G2 phase-related protein, peaked in expression at D5 and D10, lacked detectable expression from D15 and D25, and showed a modest level at D30 (daughter cells); Cyclin B1, an M-phase-related protein, exhibited maximal expression at D0, followed by a rapid reduction and near-complete absence of detectable bands from D10 to D25; Cyclin D1, a G1-to-S phase-related protein, showed decreased levels of expression at D5 and D10, high level of expression at D15 and D20, and low level expression in D25 and D30 (daughter cells) (**Fig. 2H**). Collectively, these dynamic changes in cell cycle regulatory proteins suggest that PGCCs underwent sustained endoreplication, aligning with the live cell image data in **Fig. 1C**. Western blot analysis of MET-related markers revealed a progressive downregulation of the epithelial marker Pan-CK during PGCC formation, with the most significant reduction observed at the budding phase and daughter cells (**Fig. 2I**).

Conversely, mesenchymal markers, including SNAIL, CDH2, and FN1, were robustly upregulated in PGCCs, budding PGCC populations, and their daughter cells relative to regular controls, with CDH2 and FN1 displaying the most pronounced upregulation specifically in budding phases and daughter cells (**Fig. 2I**). In addition, the marker expression depended on concentration of VCR, higher concentration of VCR showed led progressive decrease of the expression of Pan-CK but increase in the expression of of FN1 and SNAIL (D10) (**Fig. 2I**). These findings indicate that PGCCs display a more pronounced mesenchymal phenotype with high concentration of VCR compared to regular HEY cells, and the increase in the level of mesenchymal state depends on drug concentrations used for PGCC induction.

We also assessed the EMT markers in daughter cells (27^th^ generation) by Western blot and immunofluorescence analyses. The findings of Western blotting revealed that Pan-CK was down-regulated in the daughter cells, with pronounced suppression observed in daughter cells derived from 4 μM VCR-induced PGCCs (**Fig. 2J**). Meanwhile, the expression of SNAIL was up-regulated in the daughter cells of 0.5, 1 μM, as well as 4 μM VCR-induced PGCCs, while CDH2 and FN1 were increased in the daughter cells of 1 and 4 μM VCR-induced PGCCs (**Fig. 2J**).

Results of immunofluorescence staining also showed the expression of Pan-CK was reduced in the daughter cells of 4 μM VCR-induced PGCCs, while the expression of FN1 was upregulated in all daughter cells compared with regular HEY cells (**Fig. S5D**). These findings demonstrated that PGCCs exhibited significantly enhanced EMT phenotypes from the initiation of their formation compared to regular HEY cells, which were transmitted to daughter cells via budding. Consequently, daughter cells acquired more EMT characteristics relative to parental cells immediately upon formation.

The results from single PGCC-derived daughter clones showed similar results as **Fig. 2K**. Western blot analysis of epithelial (Pan-CK) and mesenchymal (FN1) markers in these clones showed a concentration-dependent downregulation of the epithelial marker Pan-CK and upregulation of the mesenchymal marker FN1 (**Fig. 2K**). These findings demonstrate that daughter cell clones derived from each single PGCC also exhibit a more aggressive phenotype compared to the regular HEY cells. Moreover, daughter cell clones from higher VCR concentrations displayed more aggressive characteristics than those induced by lower concentrations.

### 2.4 PGCCs progressively acquire enhanced stemness

To investigate the temporal dynamics of stemness in PGCCs, we used the spheroid assay to determine the stemness of PGCCs. Toward this end, we seeded 1 μM VCR-induced PGCCs at a low density (500 cells per well) into ultra-low attachment 6-well plates and cultured them in stem cell medium for 10 days. Under this condition, regular HEY cells exhibited negligible spheroid formation (>200 μm), whereas those seeded at a higher density (5,000 cells per well) demonstrated robust spheroid formation (**Fig. 3A-B**). However, in PGCCs with 500 cells per well, no spheroid-forming capacity was observed at D4 and D6, whereas a progressive increase in spheroid number was observed at D8 (**Fig. 3A-B**, > 200 μm, 18.3 ± 3.5 spheroids/well), D18 (223 ± 29.1 spheroids/well) and D21 (220 ± 20.8 spheroids/well). These findings indicate that there is a progressive activation of stemness during PGCC life cycle progression, characterized by weak stemness in the initial phase, enhanced pluripotency acquisition in the mid-phase, and peak functional stemness at the termination stage (pre-budding phase). Next, we examined the ability of the single PGCC to form spheroids by seeding 1 or 2 PGCCs/well collected at different time points into ultra-low attachment 96-well plates. Similarly, while no spheroid formation was observed in PGCCs collected at D4 and D6, spheroids began to emerge at D8 and D10, increased in frequency at D12, D15, and then peaked at D18-21 prior to budding onset (**Fig. 3C-D**).

**Figure 3.**
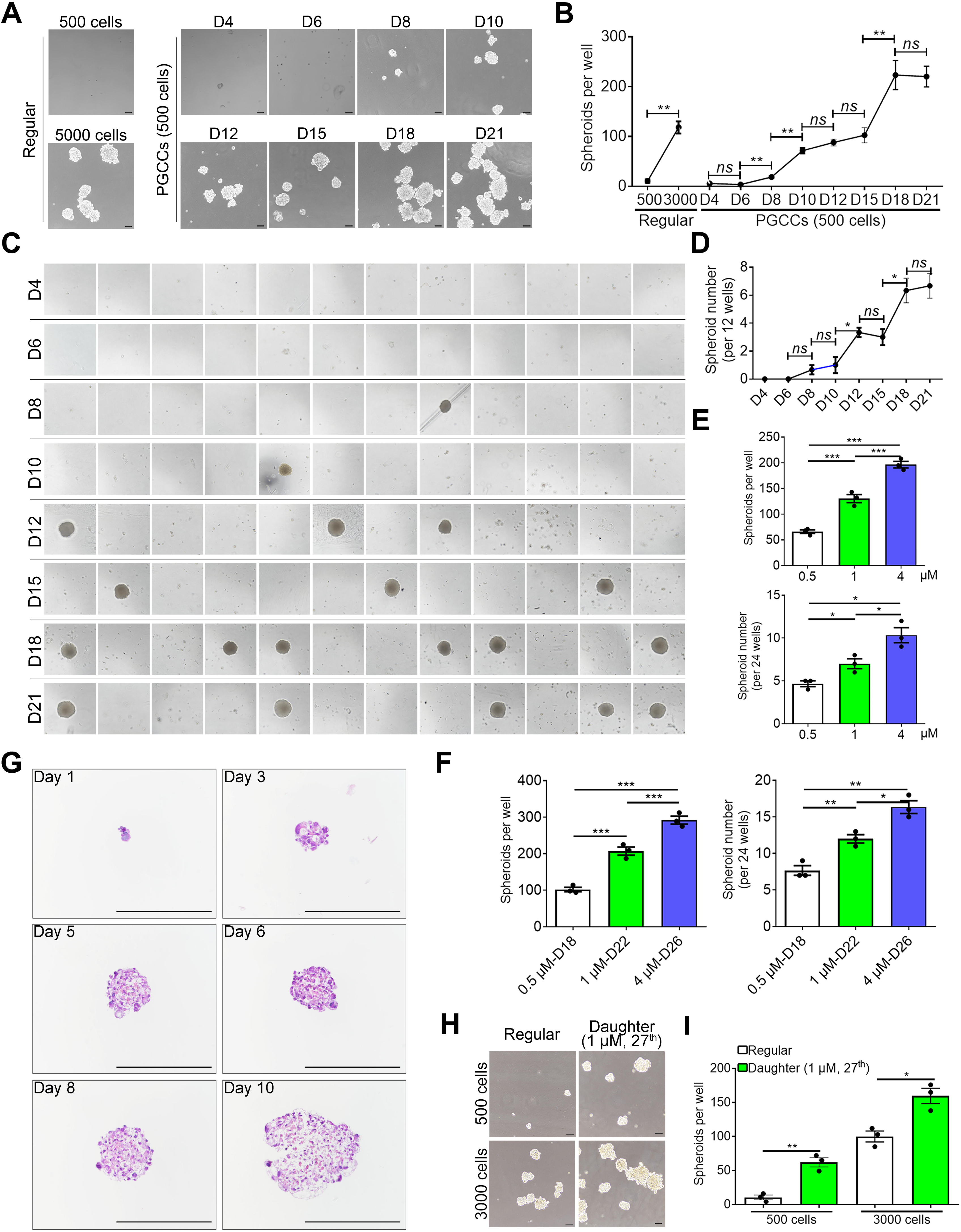
PGCCs undergo a gradual enhancement in blastomere-like stemness. **(A-B)** Regular HEY cells were seeded at 500/well or 3000/well, while PGCCs were plated at D4, 6, 8, 10, 12, 15, 18, and 21 at 500/well into ultra-low attachment 6-well plates. Spheroid formation was observed after 10 days of culture in stem cell medium. A: representative images; B: quantification of spheroid numbers per well with diameter >200 μm. Data are presented as mean ± SEM. ** *p* < 0.01. Scale bars: 200 μm. **(C-D)** PGCCs at D4/6/8/10/12/15/18/21 were seeded at 1-2 cells/well into ultra-low attachment 96-well plates. Spheroid formation was monitored after 16 days. C: Spheroid formation status per well; D: Single PGCC spheroid formation efficiency at each time point. Data are presented as mean ± SEM. *ns*, *p* > 0.05; * *p* < 0.05. Scale bars: 100 μm. **(E)** Left: PGCCs induced by 0.5, 1, and 4 μM VCR were plated at D16 into ultra-low attachment 6-well plates at a density of 500 cells/well. Spheroid formation was observed after 12 days of culture in stem cell medium. Quantification of spheroid numbers per well with diameter >200 μm. Data are presented as mean ± SEM. \*\*\**p* < 0.001. Scale bars: 200 μm. Right: PGCCs induced by 0.5, 1, and 4 μM VCR at D16 were seeded into ultra-low attachment 96-well plates at a density of 1-2 cells/well. Spheroid formation was monitored after 16 days. Single PGCC spheroid formation efficiency was analyzed. Data are presented as mean ± SEM. *p* < 0.05. Scale bars: 100 μm. **(F)** Left: PGCCs induced with 0.5, 1, or 4 μM VCR and harvested at D18, D22, or D26, respectively, were separately seeded into ultra-low attachment 6-well plates at 500 cells per well. Spheroid formation was assessed after 12-day culture in stem cell medium. Quantification of spheroid numbers per well with diameter >200 μm. Data are presented as mean ± SEM, \**p* < 0.05. Scale bars: 200 μm. Left: PGCCs induced with 0.5, 1, or 4 μM VCR and harvested at D18, D22, or D26, respectively, were separately seeded into ultra-low attachment 96-well plates at 1-2 cells per well. Spheroid formation was assessed after 16-day culture. Single PGCC spheroid formation efficiency was analyzed. Data are presented as mean ± SEM, \**p* < 0.05. Scale bars: 100 μm.(G) H&E staining of spheroids formed by 4 μM VCR-induced pre-budding PGCCs (D26) cultured in stem cell medium for days 1, 3, 5, 6, 8, and 10. **(H-I)** Daughter cells at the 27^th^ generation budded from 1 μM VCR-induced PGCCs and regular HEY cells (500 cells/well or 3000 cells/well) were cultured in ultra-low attachment 6-well plates for 14 days. E: Representative images to show spheroid formation. F: Statistical analysis of spheroids number (> 200 μm) per well in each group. Data presented as mean ± SEM. * *p* < 0.05, ** *p* < 0.01.

To investigate the differences in stemness of PGCCs induced by varying concentrations of VCR, we added PGCCs (D16) generated with 0.5, 1, and 4 μM VCR into ultra-low attachment 6-well plates (500 cells/well) and observed spheroid formation after 12 days. The results demonstrated that PGCCs induced by higher VCR concentrations formed a greater number of spheroids (>200 μm) (**upper panel of Fig. 3E, and S7A**). A single spheroid assay yielded consistent results (**lower panel of Fig. 3E, and S7B**). Additionally, when PGCCs in the pre-budding stage (generated with 0.5 μM VCR (D18), 1 μM VCR (D22), and 4 μM VCR (D26)) were separately collected for spheroid formation assays and single spheroid formation assays, similarly, the results demonstrated that PGCCs induced by higher VCR concentrations exhibited stronger spheroid-forming capacity at the pre-budding stage (**Fig. 3F and S7C-D**). Furthermore, the spheroid-forming capacity of PGCCs at this stage was uniformly superior to that of their respective D16 PGCCs (**Fig. 3E-F and Fig. S7A-D**). In summary, these findings indicate that PGCCs induced by higher VCR concentrations exhibit enhanced stemness.

Taken together, formation of PGCCs is associated with decreased proliferation, while budded daughter cells gradually resume the proliferative activity (**Fig. S8**). Formation of PGCCs and budded daughter cells is associated with increase in stemness (**Fig. S8**). For EMT, the process undergoes progressive upregulation from parental cells and through the life cycle of PGCCs and budded daughter cells (**Fig. S8**).

### 2.5. PGCC-derived spheroids are capable of differentiation into three-germ-layer-derived tissue structures or cell lineages

We simultaneously performed H&E staining on the spheroids formed by culturing 4 μM VCR-induced pre-budding PGCCs (D26) in stem cell medium on days 1, 3, 5, 6, 8, and 10. It was observed that as the spheroids increased in size, the cells exhibited an increased level of cellular heterogeneity but remain in undifferentiated state, consistent with the strong stemness of PGCCs (**Fig. 3G**).

Additionally, daughter cells derived from 1 μM VCR-induced PGCCs (27^th^ generation) exhibited enhanced spheroid formation capacity relative to regular cancer cells when plated into ultra-low attachment 6-well plates at either 500 cells/well (*p* < 0.05) or 3000 cells/well (*p* < 0.05) densities, though they formed fewer spheroids than those of PGCCs at D21 (**Fig. 3H-I**). These data highlight progressive stem-like property acquisition during PGCC formation and maintenance, peaking before the budding stage.

To investigate the embryonic differentiation potential and tissue regenerative capacity of PGCCs, we seeded pre-budding PGCCs (500 cells per well) and regular HEY cells (5,000 cells per well) onto ultra-low-attachment six-well plates and cultured them in stem cell medium for 3 days. The medium was subsequently replaced with STEMdiff™ Trilineage medium and maintained for additional periods of an additional 1, 3, 5, and 10 days (modified endoderm condition), 5 days (mesoderm condition) and 7 days (ectoderm condition). We observed that spheroid formation occurred only in the modified endoderm medium in both regular HEY cells and PGCC cultures on day 5 (**Fig. 4A**). However, spheroids derived from regular HEY cells progressively lost viability after 10 days of culture. In contrast, PGCC-derived spheroids on day 5 exhibited progressive increases in cell size accompanied by marked structural remodeling. Formalin fixation followed by H&E staining revealed that PGCC-derived spheroids on day 5 developed lumen-like structures resembling organoid architecture (**Fig. 4A**, lower panels). Notably, lumen formation was more complete at higher vincristine (VCR) concentrations (**Fig. 4A**, lower panels).

**Figure 4.**
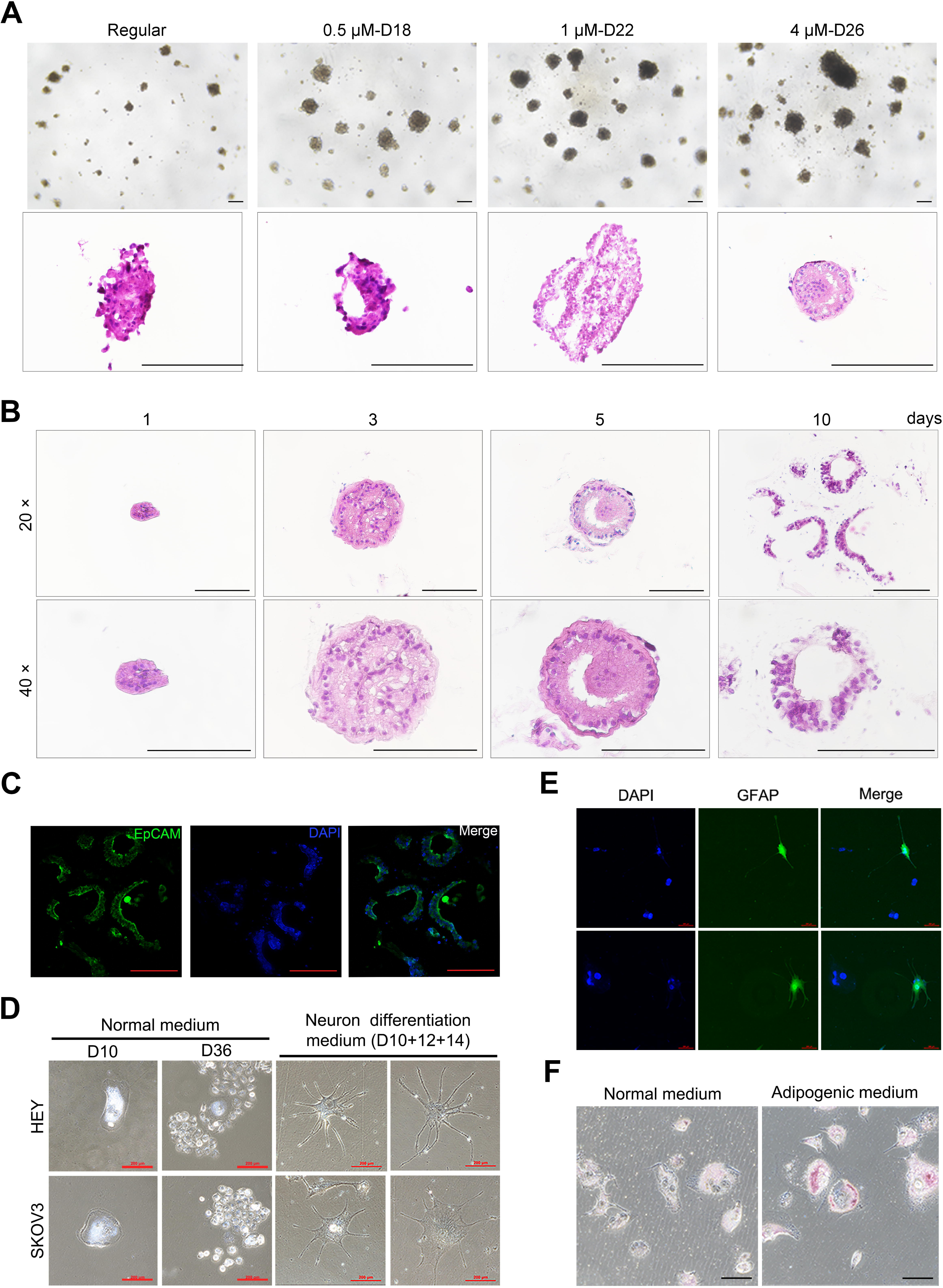
PGCCs acquire multipotent differentiation potential in a concentration-dependent manner. **(A)** Embryonic differentiation potential of PGCCs induced with 0.5, 1, or 4 μM VCR. Spheroid formation assay of pre-budding stage PGCCs (500 cells/well) and regular HEY cells (5,000 cells/well) cultured in ultra-low attachment 6-well plates. Cells were initially maintained in stem cell medium for 3 days, then switched to STEMdiff™ Endoderm Media for 5 days. Upper layer: Bright fields. Lower layer: H&E staining. Scale bars: 200 μm. **(B)** PGCCs induced by 4 μM VCR were cultured in STEMdiff™ Endoderm Media for 1, 3, 5, and 10 days, and spheroids were separately collected at each time point for H&E staining. Scale bars: 200 μm. **(C)** After culturing PGCCs (induced by 4 μM VCR) in STEMdiff™ Endoderm Media for 10 days, spheroids were collected for EpCAM staining. Scale bars: 100 μm. **(D-E)** Morphological characterization of PGCCs (D10 induced with 1 μM VCR; 3000 cells/well in 6-well plates) following 12 days of initial culture in STEMdiff^TM^ Forebrain Neuron Differentiation Medium and subsequent 14-day culture in maturation medium. Left panel of J: Bright-field microscopy images of HEY-derived PGCCs at D10 and D36. Right panel of J: PGCCs cultured with neuron differentiation and maturation medium. Scale bars: 200 μm. K: Immunofluorescence staining of (GFAP) in STEMdiff^TM^-differentiated PGCCs: Scale bars: 200 μm. **(F)** HEY-derived PGCCs (D10) were induced to differentiate into adipocyte-like cells. After differentiation, intracellular lipid was stained with Oil Red O. Upper panel: PGCCs cultured with normal culture medium; Lower panel: PGCCs cultured with adipogenesis differentiation medium.

Temporal H&E analysis of spheroids collected at multiple time points (derived from pre-budding PGCCs (D26) induced by 4 μM VCR) demonstrated that lumen structures became progressively more organized and morphologically mature with prolonged culture (**Fig. 4B**). In one spheroid, we observed a fully developed fibrous capsule surrounding a morphologically mature epithelial lining accompanied by stromal and vascular-like components (**Fig. S7E**).

Additional spheroids stratified by size and differentiation stage (day 1, 3, 5, and 10) were shown in **Fig. S9**, illustrating the stepwise developmental progression toward structured tissue architecture. Notably, thick fibrous capsules were observed at the periphery of immature lumen-forming spheroids (**Fig. S9, left lower panels**). These observations suggested that PGCC nuclei initially underwent marked nuclear expansion, followed by radial self-organization into spatially distinct peripheral and central nuclear territories. The peripheral nuclear cohort survived and diverged into two major lineages: a subset adopted fibroblast-like features contributing to fibrous capsule formation, whereas others underwent epithelialization and further self-organization to form the luminal epithelial wall. In contrast, the central nuclear cohort underwent programmed clearance, generating a proto-lumen with an initially irregular luminal surface that subsequently matured into a structurally stabilized glandular architecture (**Fig. 4A–B** and **Fig. S9**).

Consistent with these in vitro findings, we also identified central clearance within lumen-forming regions in multiple human tumor specimens, including high-grade serous carcinoma (**Fig. S10A**), endometrial hyperplasia (**Fig. S10B**), and endometrioid adenocarcinoma (**Fig. S10C–D**). Immunofluorescence analysis further demonstrated uniform EpCAM expression in the lumen-lining cells, confirming their epithelial identity (**Fig. 4C**).

Although PGCCs failed to form spheroids in both mesoderm and ectoderm media, we nevertheless sought to investigate the lineage-specific differentiation of PGCCs associated with the mesoderm and ectoderm. Considering that the ectoderm is primarily linked to neural differentiation, we sub-cultured the PGCCs in neuron differentiation and maturation medium with concurrent SMAD signaling inhibition throughout the differentiation and maturation processes. PGCCs exhibited neuron-like morphology when cultured in differentiation-inducing medium, whereas no such morphological change was observed in PGCCs maintained in regular control medium (**Fig. 4D**). Immunofluorescence staining results showed that these cells were positive for GFAP and negative for TUBB3 (**Fig. 4E** and **S11B**). In contrast, regular cancer cells exhibited negative immunostaining for both GFAP and TUBB3 (**Fig. S11A**), suggesting these PGCCs have developed astrocyte-like characteristics. Furthermore, given that the mesoderm is associated with adipogenic differentiation, we induced adipocyte differentiation in PGCCs (D10) using adipogenic differentiation medium. The results demonstrated that this medium successfully promoted PGCCs to differentiate into adipocytes, consistent with prior reports (**Fig. 4F**) 24,29. Collectively, these data reveal that PGCCs undergo a progressive enhancement in blastomere-like stemness.

### 2.6 PGCCs partially exhibited senescent phenotypes and showed increased expression of SASP markers

To examine if VCR-induced PGCCs are associated with a senescence phenotype, SA-β-gal staining was performed on PGCCs obtained from HEY and SKOV3 cells treated with 0.5, 1, and 4 μM of VCR at D10. Consistent with our early observations in paclitaxel ^24^ and olaparib-induced PGCCs ^13^, HEY and SKOV3-derived PGCCs at D10 induced by VCR exhibited enhanced SA-β-gal activity compared with regular cancer cells (**Fig. 5A**). Western blot analysis of two known senescence regulators p16 and phosphorylated Rb (p-Rb) showed increased expression of p16 and decreased p-Rb protein levels in PGCCs at D10 compared with regular HEY and SKOV3 cells, respectively, supporting the activation of the senescence pathway in PGCCs (**Fig. 5B**).

**Figure 5.**
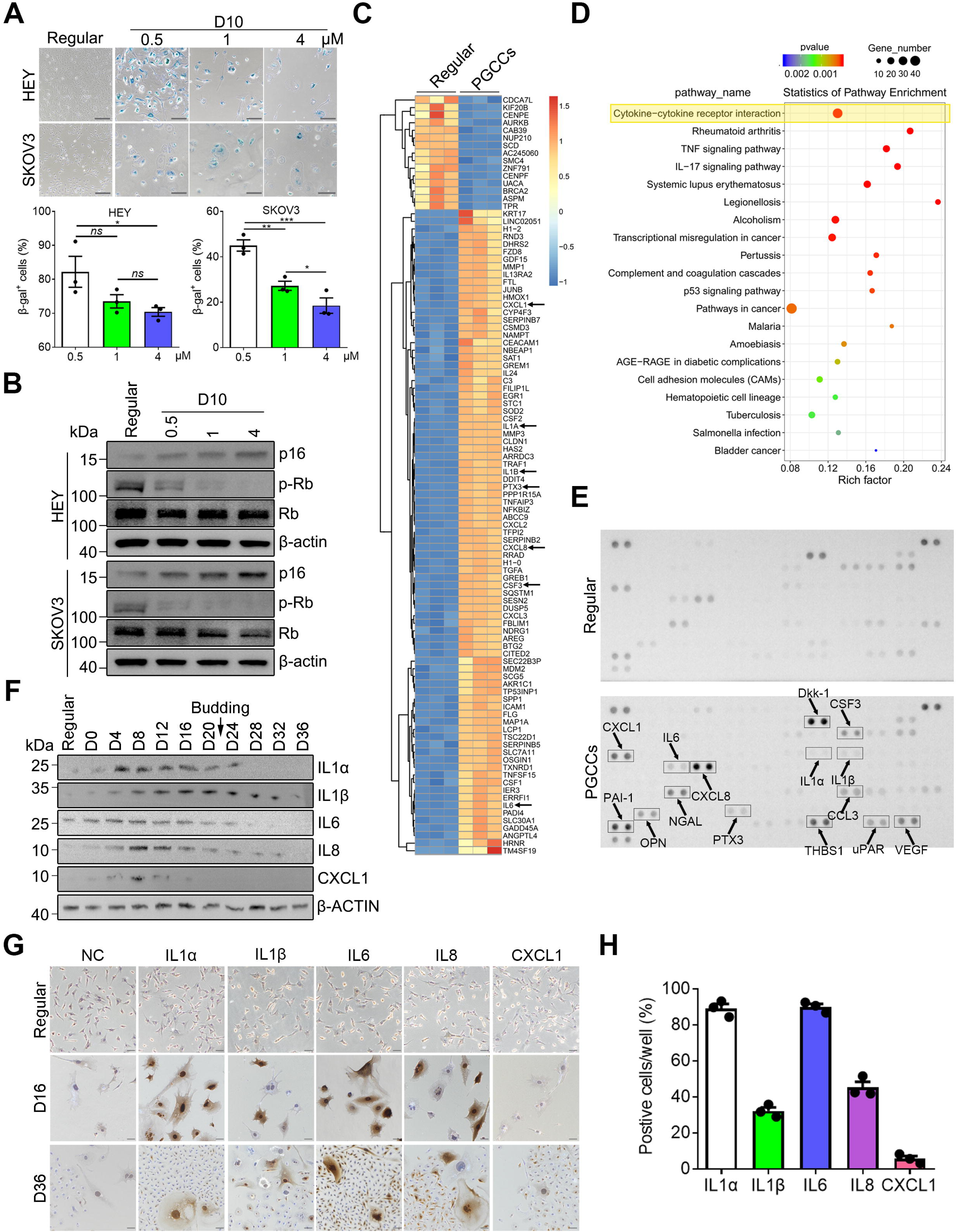
VCR-induced PGCCs partially exhibit senescent phenotypes and SASP. **(A)** HEY and SKOV3 cells were treated with varying concentrations of VCR for 2 days, followed by a 10-day recovery period in drug-free medium. PGCCs were then stained with SA-β-gal. Scale bars: 200 μm. Upper panel: representative images; lower panel: Quantification of SA-β-gal-positive (SA-β-gal) PGCCs across groups. Data are presented as mean ± SEM. *ns*, *p* > 0.05, * *p* < 0.05, ** *p* < 0.01, *** *p* < 0.001. **(B)** Western blot analysis of p16, p-Rb, and total Rb protein levels in PGCCs (D10; induced by different VCR concentrations) and regular cancer cells. **(C)** Heatmap visualization of differentially expressed genes (DEGs) identified by RNA sequencing, comparing VCR-induced PGCCs (1 μM, D10) with regular HEY cells. **(D)** KEGG pathway enrichment analysis of DEGs, with the cytokine-cytokine receptor interaction pathway highlighted as the most significantly enriched. **(E)** Proteome Profiler^TM^ Human Cytokine XL Array analysis of supernatants from VCR-induced PGCCs (D10) and regular HEY cells. Upregulated proteins in PGCC supernatants are labeled. **(F)** Western blotting to detect the protein levels of IL1α, IL1β, IL6, IL8, and CXCL1 in PGCCs induced by 1 μM of VCR-treated HEY cells at D0, D4, D8, D16, D20, D24, D28, D28, and D36 compared with regular HEY cells. **(G-H)** PGCCs at D16 and D36 and regular HEY cells were subjected to ICC assays to determine the expression of IL1α, IL1β, IL6, IL8, and CXCL1 during PGCC formation and budding. Scale bars: 100 μm. H: The percentages of positive-stained IL1α, IL1β, IL6, IL8 and CXCL1 in PGCCs at D16 were analyzed, respectively. Data were presented as mean ± SEM.

To determine the change of mRNA expression levels during the life cycle of PGCCs, we performed the RNA sequencing (RNA-seq). As shown in **Fig. 5C**, the RNA-seq results revealed significant transcriptomic changes during the PGCC formation. Compared with the regular HEY cells, there was a massive upregulation of genes in PGCCs involved in cytokine-cytokine receptor interaction (**Fig. 5D**). Integration of RNA-seq data with human XL cytokine array data (**Fig. 5E**) revealed a subset of cytokines up-regulated in PGCCs that were correlated with the senescence-associated secretory phenotype (SASP), including IL1α, IL1β, IL6, CXCL1, IL8, CSF3 and PTX3. These results were further validated by qRT-PCR (**Fig. S12A-B**). To further elucidate their potential roles in PGCCs, we performed systematic temporal profiling of these SASP-associated proteins across PGCCs’ life cycle using Western blot analysis. As demonstrated in **Fig. 5F**, IL1α, IL1β, IL6, IL8 and CXCL1 exhibited upregulated protein expression during early PGCC formation relative to regular HEY cells, with levels returning to baseline in late PGCC stages and in daughter cells. CSF3/PTX3 remained undetectable throughout all experimental time points. These findings were corroborated by ELISA (**Fig. S12C**) and immunocytochemistry (ICC) analyses (**Fig. 5G-H**). Interestingly, ICC analysis of PGCCs at D16 showed that the majority of PGCCs exhibited positive expression of IL1α (89.1 ± 4.4%) and IL6 (89.9 ± 2.9%), yet only a subset of PGCCs showed positive expression of IL1β (32.1 ± 3.4%) and IL8 (45.3 ± 5.4%). In contrast, CXCL1 positivity was found in only very few PGCCs (5.64 ± 2.6%), with most remaining predominantly negative (**Fig. 5G-H**). These results demonstrate significant heterogeneity in chemokine and cytokine expression profiles among PGCCs.

### 2.7 Inhibition of IL1**β**, IL6, or IL8 pathways impairs the formation, budding capacity, stemness, and differentiation potential of PGCCs

To determine the role of these SASP-associated factors in the fate and function of PGCCs, siRNAs were transfected into HEY cells for 2 days to silence the expression of IL1α, IL1β, IL6, IL8 and CXCL1 (**Fig. S13**), respectively. PGCC formation was induced by treating cells with 1 μM VCR for 2 additional days, followed by recovery in drug-free culture medium. At D20, we observed that knockdown of IL1α or CXCL1 in HEY cells did not impair their capacity to form PGCCs (**Fig. 6A-B**). In contrast, knockdown of IL1β (*p* < 0.05), IL6 (*p* < 0.05) or IL8 (*p* < 0.01) significantly reduced VCR-induced PGCC formation, with particularly pronounced effects in cells with IL1β or IL8 knockdown (**Fig. 6A-B**). At D30, PGCCs in the IL1α, IL6, and CXCL1 knockdown groups retained the ability to generate daughter cells. In contrast, those in the si-IL1β and si-IL8 groups failed to produce viable progeny (**Fig. 6A**). When we specifically antagonized IL1R1, IL-6R, and CXCR1/CXCR2 in HEY and SKOV3 cells with AF12198 (10 nM), tocilizumab (1 μM), and navarixin (1 μM), respectively, similar results were obtained (**Fig. S14A-C**).

**Figure 6.**
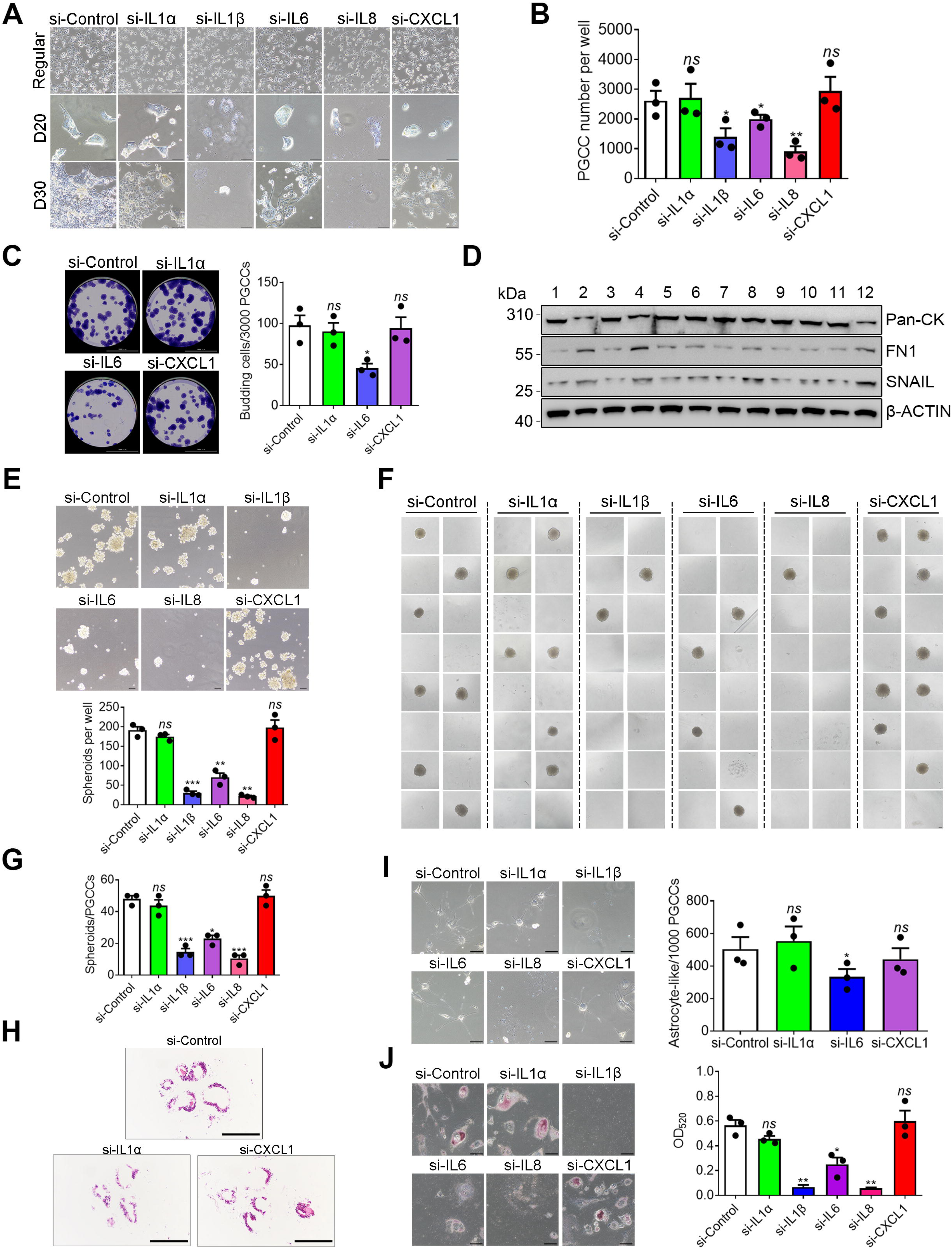
Knockdown of IL1β, IL6 as well as IL8 impairs PGCC formation, budding, and multipotent differentiation potential. **(A-B)** IL1α, IL1β, IL6, IL8, or CXCL1 was knocked down in HEY cells, respectively. Then, 1 μM of VCR was added for 2 days and allowed to recover in a drug-free medium to induce the formation of PGCCs. A: Morphological analysis was performed at D20 and D30. Scale bars: 100 μm. B: Quantification of PGCC numbers per well (D20) across knockdown groups. **(C)** VCR-induced PGCCs (3000 per well) at D10 from HEY with IL1α, IL6, or CXCL1 silencing were subcultured into 12-well plates and maintained for 30 days. Crystal violet was used to stain the budding clones. Scale bars: 10 mm. Left panel: Representative images; Lower panel: Quantification of clone numbers. ns, *p* > 0.05, * *p* < 0.05. **(D)** Western blotting to detect the protein levels of EMT markers Pan-CK, FN1, and SNAIL in PGCCs and regular HEY cells in IL1α, IL1β, IL6, IL8, or CXCL1 knockdown groups. 1: regular HEY cells with si-Control; 2: PGCCs with si-Control; 3: regular cells with si-IL1α; 4: PGCCs with si-IL1α; 5: regular cells with si-IL1β; 6: PGCCs with si-IL1β; 7: regular cells with si-IL6; 8: PGCCs with si-IL6; 9: regular cells with si-IL8; 10: PGCCs with IL8; 9: regular cells with si-CXCL1; 10: PGCCs with CXCL1. **(E)** PGCCs at D20 from IL1α, IL1β, IL6, IL8, or CXCL1 knockdown groups were cultured in ultra-low attachment 6-well plates (500 cells/well) for 10 days using stem cell medium. Upper panel: Representative spheroid formation images; Lower panel: Quantification of spheroids (>200 μm diameter). Scale bars: 200 μm. Data are presented as mean ± SEM. *ns*, *p* > 0.05; **, *p* < 0.01; ***, *p* < 0.001 (vs. control). **(F-G)** Single PGCCs isolated at D20 from IL1α, IL1β, IL6, IL8, or CXCL1 knockdown groups were cultured in ultra-low-attachment 96-well plates at a density of 1-2 cells/well using a defined stem cell medium for 16 days. F: Spheroid formation status visualized under phase-contrast microscopy. G: Quantitative analysis of spheroid formation efficiency of single PGCCs across control and knockdown groups. Scale bars: 100 μm.. Data are presented as mean ± SEM. *ns*, *p* > 0.05; *, *p* < 0.05; ***, *p* < 0.001 (*vs*. control). **(H)** Knockdown of IL1α, IL1β, IL6, IL8, and CXCL1 was performed in regular HEY cells. Subsequently, 4 μM VCR was added for 2 days, followed by a recovery period to induce PGCC formation. Twenty-six days post-induction, these PGCCs were cultured in STEMdiff™ Endoderm Media for 10 days, and spheroids were harvested for H&E staining. (I)Following knockdown of IL1α, IL1β, IL6, IL8, and CXCL1 in regular HEY cells, 1 μM VCR was added for 2 days with recovery to induce PGCC formation, and ten days later cells were cultured in STEMdiff™ forebrain neuron differentiation medium (12 days) and maturation medium (14 days, 0.5 μM of LDN-193189 and 10 μM of SB431542 were added). Left panel: Representative cell morphology was observed in the bright field. Scale bars: 200 μm. Right panel: The numbers of astrocyte-like cells in each well were calculated. Data are presented as mean ± SEM. *ns*, *p* > 0.05, * *p* < 0.05. **(J)** IL1α, IL1β, IL6, IL8, and CXCL1 were knocked down in regular HEY cells, then 1 μM of VCR was added for 2 days and allowed for recovery to induce the formation of PGCCs. Ten days later, the cells were cultured were cultured with Adipogenic Differentiation A medium for 2 days and Adipogenic Differentiation A medium for 1 day, repeated three times. Left panel: Representative cell morphology was observed in the bright field. Scale bars: 200 μm. Right panel: To quantify the amount of stained Oil Red O, the dye was eluted with isopropyl alcohol, and its optical density was monitored spectrophotometrically at 520 nm. Data are presented as mean ± SEM. *ns*, *p* > 0.05, * *p* < 0.05, ** *p* < 0.01.

To further compare the budding capacity of the PGCCs in the IL1α, IL6, and CXCL1 knockdown groups, we subcultured the PGCCs at D10 into 12-well plates at a density of 3000 cells per well and maintained them for 30 days. Crystal violet staining results revealed that only IL6 knockdown (*p* < 0.05) significantly decreased the budding percentage of PGCCs (**Fig. 6C**). Inhibiting IL6R with tocilizumab also suppressed the budding capacity of PGCCs (**Fig. S14D**). Meanwhile, we examined the expression changes of the EMT markers in these PGCCs and their corresponding regular cancer cells. The results demonstrated that both IL1β and IL8 knockdown attenuated the reduced expression trend of the epithelial marker Pan-CK and suppressed the upregulation trends of mesenchymal markers FN1 and SNAIL during PGCC formation (**Fig. 6D**). In contrast, IL1α and CXCL1 knockdown exhibited no significant effects, while IL6 showed intermediate efficacy in modulating these EMT-related markers (**Fig. 6D**). Furthermore, the spheroid formation capacity of PGCCs (**Fig. 6E**) and single PGCCs (**Fig. 6F-G**) at D20 in the IL1β, IL6 as well as IL8 knockdown groups was significantly impaired compared to the control group, with the most pronounced reduction observed in the IL-1β and IL-8 knockdown groups.

Finally, we subjected these PGCCs to endoderm differentiation culture under identical conditions.

The results showed that after 10 days, only PGCCs from the control group and those with IL1α or CXCL1 knockdown formed lumen structures, whereas PGCCs with IL1β, IL6, or IL8 knockdown failed to form spheroids (**Fig. 6H**). Additionally, we also cultured these PGCCs at D10 using neuron differentiation and maturation medium and adipose differentiation medium.

The results showed no viable astrocyte-like cells (**Fig. 6I**) and adipocyte-like cells (**Fig. 6J**) in the si-IL1β and si-IL8 groups. Notably, IL6 knockdown markedly impaired PGCC differentiation compared to si-Control, si-IL1α, and si-CXCL1 groups (**Fig. 6I-J**). Meanwhile, we demonstrated that antagonizing IL1R1, IL-6R, and CXCR1/CXCR2 in HEY and SKOV3 cells using AF12198, tocilizumab and navarixin, respectively, each potently inhibited the induction of PGCCs into adipocyte-like cells (**Fig. S14E**). Collectively, these findings suggest that IL1β, IL6, and IL8 play indispensable roles in governing the fate of PGCCs, with IL1β and IL8 exhibiting dominant functional roles in driving their phenotypic plasticity and stemness.

## 3 Discussion

A central innovation of this study is the systematic delineation of the spatiotemporal dynamics underlying the formation of architecturally defined glandular tissue derived from conventional tissue-cultured cancer cells, in addition to the generation of three-germ-layer differentiation phenotypes and therapy-resistant progeny. We demonstrate that the fate transition of cancer cells toward endoderm-derived, mature structured tissue is strongly dependent on the dose of chemotherapeutic stress. These findings reveal that cancer cells are highly plastic and capable of context-dependent reprogramming in response to stress intensity, thereby supporting our earlier hypothesis that the PGCC life cycle recapitulates a blastomere-like pre-embryonic program ^24^.

At the tissue level, we provide a spatiotemporal dynamic view of tissue formation through reactivation of the PGCC life cycle in somatic cancer cells, highlighting the unique capacity of PGCCs for higher-order structural tissue generation (Fig. 4). Notably, PGCC-derived spheroids form lumen-like structures that are morphologically and developmentally distinct from those formed by regular HEY cells (Fig. 4A-B). Furthermore, under defined conditions, PGCCs exhibit multilineage differentiation potential, giving rise to astrocyte-like cells (ectodermal lineage; Fig. 4D–E) and adipocyte-like cells (mesodermal lineage; Fig. 4F). Collectively, these observations indicate reactivation of blastomere-like developmental programs that confer embryonic-level potency on PGCCs.

Using a gradient of vincristine (VCR) concentrations, we further demonstrate that both glandular structures and daughter cells derived from PGCCs exhibit dose-dependent behavioral divergence.

Increasing drug intensity results in a progressive decline in proliferation of early-budded daughter cells (Fig. 2B, S5A), consistent with our prior report. The initially budded progeny from mother PGCCs are highly heterogeneous; however, with continued passaging, highly proliferative clones gradually emerge through clonal expansion, ultimately surpassing regular cancer cells and acquiring a more aggressive phenotype (Fig. 2B, S5A).

Polyploid cells have long been recognized for their capacity for dedifferentiation and multidirectional differentiation and are frequently associated with cellular senescence and the senescence-associated secretory phenotype (SASP), a complex network of proinflammatory cytokines, chemokines, and growth factors. However, how SASP paradoxically promotes PGCC survival and plasticity has remained unclear. Our data support a model in which SASP produced by senescent PGCCs enhances regenerative potential through coordinated autocrine and paracrine signaling. Specifically, IL-1β, IL-6, and IL-8 play pivotal roles in PGCC formation, self-renewal, and tissue morphogenesis via activation of blastomere-like programs, positioning SASP cytokines as master regulators of PGCC plasticity and tissue genesis.

Collectively, our findings support a unifying model in which chemotherapy-induced polyploidization of senescent cancer cells drives not only therapy resistance but also structured tissue regeneration. As summarized in Fig. 7, the PGCC life cycle comprises four interlinked phases—initiation, self-renewal, termination, and stability—each governed by coordinated stress and developmental signaling pathways that collectively regulate progeny regeneration and tissue remodeling.

**Figure 7.**
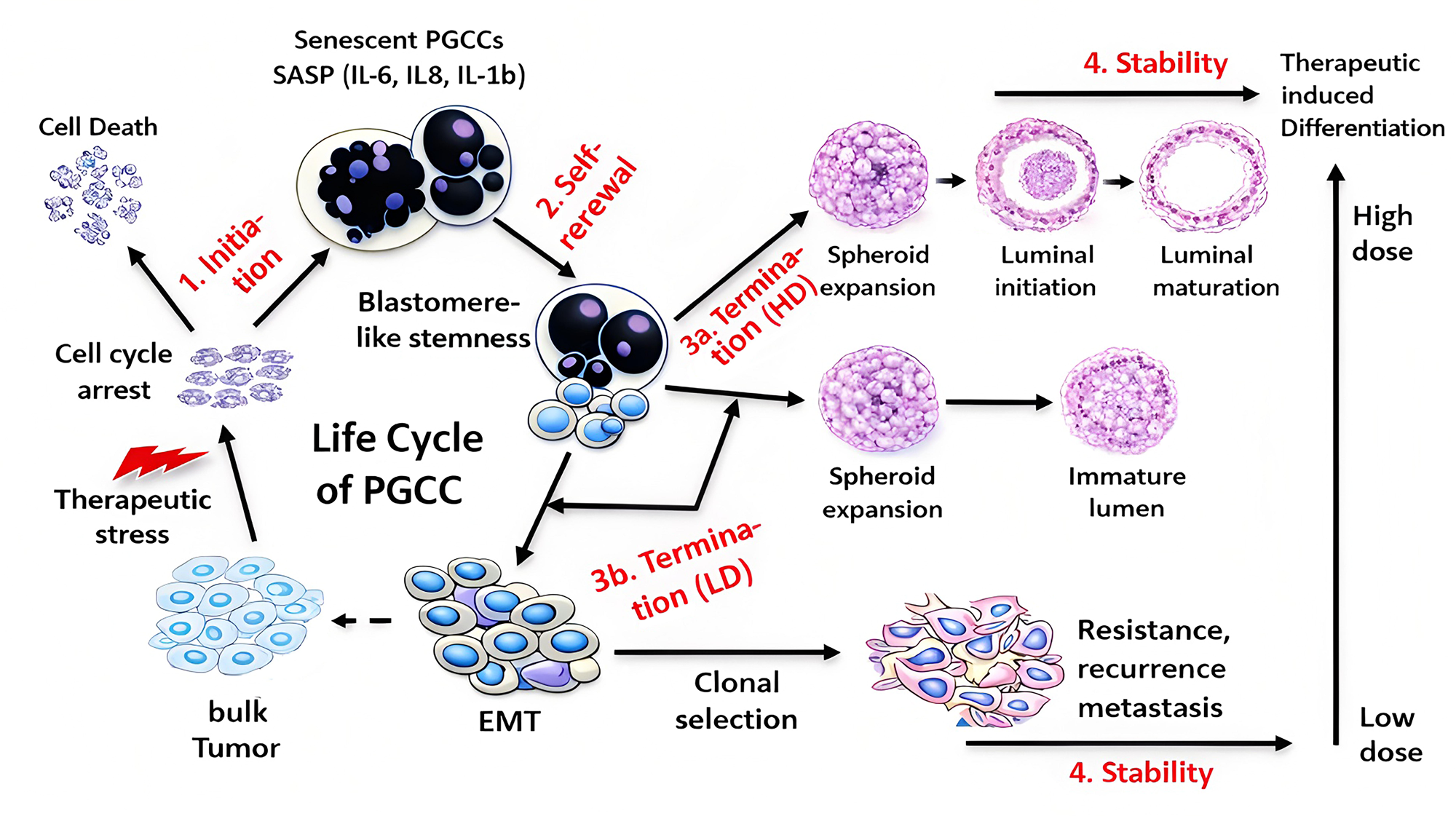
Schematic representation of the complete fate trajectory of life cycle of polyploid giant cancer cell (PGCC) induced by chemotherapeutic drugs. Chemotherapyinduces tumor cell reprogramming through four interconnected stages: (1) Initiation: begins with therapy-induced mitotic suppression, forcing endoreplication in bulk cancer cells. While most proliferative cells undergo apoptosis, a senescent subpopulation survives via genome amplification, generating PGCCs marked by SASP cytokines (IL6/8/1β). (2) Self-Renewal: where PGCCs activate EMT pathways and SASP, reprogramming into blastomere-like stemness resembling early embryonic states. After a latent period, these reprogrammed PGCCs transition into (3) Termination: diverging based on dose: high-dose (3a) stabilizes PGCCs in a dormant state, uniquely triggering tissue-differentiation; low-dose (3b) induces undifferentiated/poorly differentiated spheroids that bud EMT-driven, heterogeneous drug-resistant progeny fueling recurrence/metastasis; (4) Stability: high dose results in therapy-induced differentiation into mature glandular tissue from immature proto-lumen into matured tissue structure, while low dose results in clonal selection establishes genetically/phenotypically stable, treatment-refractory populations for recurrence and metastasis. Dashed arrow indicate the budded daughter cells can be back to the life cycle of PGCCs in response to new stress.

Following both low- and high-dose chemotherapy, the initiation and self-renewal stages are largely similar:

### (1) Initiation

This phase is triggered by therapy-induced mitotic suppression, during which the majority of proliferating tumor cells undergo apoptosis, whereas a small subpopulation enters senescence and undergoes endoreplication to generate PGCCs. This represents an adaptive survival entry point in response to catastrophic stress, enabling escape from mitotic catastrophe through whole-genome amplification and genomic resetting.

### (2) Self-renewal

During this phase, PGCCs progressively acquire EMT features and activate SASP cytokines (IL-1β, IL-6, IL-8), which together drive reprogramming toward blastomere-like stemness reminiscent of early embryonic stages. After a latency of approximately 2–3 weeks, reprogrammed PGCCs enter the termination phase.

### (3) Termination

At this stage, cell fate diverges in a dose-dependent manner.

Under low-dose conditions, spheroids remain undifferentiated or poorly differentiated, maintaining a high nuclear-to-cytoplasmic ratio and budding highly heterogeneous daughter cells that subsequently undergo EMT to generate proliferative progeny.

In contrast, high-dose–induced PGCCs undergo a prolonged termination phase characterized by structural morphogenesis rather than progeny budding. During spheroid growth, PGCC nuclei undergo radial territorial self-organization, establishing spatially distinct peripheral (productive) and central (sacrificial) nuclear domains. This architecture primes the cell for structured tissue morphogenesis.

### (4) Stability

Under low-dose conditions, spheroids ultimately disintegrate into heterogeneous, partially reprogrammed cells that acquire proliferative capacity and generate resistant progeny responsible for recurrence, metastasis, and long-term tumor persistence.

Under high-dose conditions, however, the peripheral nuclear cohort integrates with the fibrous capsule, acquires epithelial polarity, and self-organizes into mature epithelial structures, while the central nuclear cohort is selectively eliminated to generate an incipient lumen for lineage growth and expansion. Importantly, this process represents programmed partial nuclear sacrifice rather than global cell death, enabling organized glandular morphogenesis and expansion.

Because glandular architecture represents the structural foundation of many major organs—including pancreas, stomach, intestine, breast, prostate, and endometrium—dysregulation of this developmental program may contribute directly to tumorigenesis in addition to therapeutic resistance. Elucidating the mechanisms governing therapy-induced glandular morphogenesis therefore has broad implications for understanding both tumor tissue remodeling and normal organ development.

Our findings also have important therapeutic implications. Tumors from different patients exhibit heterogeneous sensitivities to therapy due to diverse genetic backgrounds, yet current treatment paradigms largely apply uniform dosing. Our data suggest that dose intensity critically influences PGCC fate decisions, potentially contributing to variable resistance outcomes observed clinically. These observations support a future precision strategy incorporating personalized stress dosing and raise the possibility that promoting therapy-induced maturation toward benign, structured tissue states may represent a novel avenue to limit resistance and disease progression.

## Materials and methods

### 4.1 Cell culture and drugs

The ovarian cancer cell lines HEY (RRID: CVCL_0297) and SKOV3 (RRID: CVCL_0532) were obtained and cultured as previously described by our laboratory ^30^. Briefly, HEY cells were cultured in RPMI 1640 medium (Biological Industries (BI), Beit Haemek, Israel) supplemented with 10% (v/v) fetal bovine serum (FBS; Gibco, Rockville, MD), 100 U/mL penicillin (Vivacell, Shanghai, China), and 100 μg/mL streptomycin (Vivacell). SKOV3 cells were maintained in McCoy’s 5A medium (BI) with same additives. All cell lines were incubated at 37°C in a humidified atmosphere containing 5% CO_2_. All cell lines employed in this study have undergone Short Tandem Repeat (STR) authentication to confirm their identities. The VCR sulfate (HY-N0488), AF12198 (HY-P1110), tocilizumab (HY-P9917) and navarixin (HY-10198) were purchased from MedChemExpress (MCE, Shanghai, China).

### 4.2 Cell cycle analysis

Treated HEY and SKOV3 cells were harvested, washed three times with phosphate-buffered saline (PBS; BI), and fixed overnight at -20°C in 1 mL of 70% ethanol (Lingfeng Chemical, Shanghai, China). Before flow cytometric analysis, cells were centrifuged to pellet, resuspended in PBS, and washed again. The cell pellet was then resuspended in PBS containing 0.05 mg/mL propidium iodide (PI; Beyotime, Shanghai, China) and 1 mg/mL RNase A (Beyotime), followed by incubation at 37°C for 30 minutes. Finally, samples were analyzed using a BD FACSCalibur flow cytometer (RRID: SCR_000401; BD Biosciences, Franklin Lakes, NJ), and the data were processed using FlowJo software (RRID: SCR_008520; Tree Star, Eugene, OR).

### 4.3 Colony formation assay

To detect the budding percentage of PGCCs, HEY and SKOV3 cells were treated with varying concentrations of VCR for 2 days and allowed to recover in culture for 10 days. Resulting PGCCs (HEY: 3000 cells/well; SKOV3: 1500 cells/well) were then collected and subcultured into 12-well plates at the cell density specified in the results. After a subsequent 30-day recovery period, the cells were fixed with 4% paraformaldehyde (Sangon, Shanghai, China) for 30 minutes, air-dried, and stained with 0.05% crystal violet (Beyotime) for 15 minutes. The plates were gently washed with double-distilled water (ddH_2_O) and air-dried again. Finally, colony formation was quantified by manual counting, and representative images were captured using a Lionheart FX automated microscope (RRID: SCR_019744; Agilent Technologies, Santa Clara, CA). To assess the colony-forming capacity of regular HEY and the daughter cells of PGCCs, 2000 cells/well were seeded in 12-well plates and maintained under standard culture conditions for 14 days before colony quantification.

### 4.4 SRB assay

The PGCC-derived daughter cells of different generations and regular HEY cells were seeded into 24-well plates at a density of 5 × 10^4^ cells per well in RPMI 1640 medium supplemented with 5% (v/v) FBS. The culture media were refreshed every 48 hours. At 0, 1, 2, and 3 days post-seeding, the cells in each well were fixed with 10% (w/v) trichloroacetic acid (TCA), stained with 0.4% (w/v) sulforhodamine B (SRB; Sigma-Aldrich, Shanghai, China), washed twice with 1% (v/v) acetic acid, and then measured for optical density (OD) at 515 nm using a BMG LABTECH POLARstar Omega microplate reader (RRID: SCR_025024; BMG LABTECH GmbH, Ortenberg, Germany).

### 4.5 MTS analysis

The PGCC-derived daughter cells of different generations and regular HEY cells were seeded into 96-well plates at a density of 1 × 10^4^ cells per well in RPMI 1640 medium supplemented with 5% FBS. The culture media were refreshed every 24 hours. Three days later, the cells in each well were added 20 µL MTS working solution (Promega, Madison, WI), incubated at 37°C for 3 h, and then measured for optical density (OD) at 490 nm using a BMG LABTECH POLARstar Omega microplate reader.

### 4.6 Western blotting

Western blotting was performed following previously established procedures ^17^. The primary antibodies are listed in **Table S1**.

### 4.7 Cell labeling and time-lapse imaging

The FUCCI (Fluorescence Ubiquitination-based Cell Cycle Indicator) system employs a red fluorescent protein (RFP) fused to the cell cycle regulator Cdt1 and a green fluorescent protein (GFP) fused to the cell cycle regulator geminin to visualize cell cycle phases. HEY cells were double-labeled with pmEGFP-α-tubulin-C1 (RRID: Addgene_21042; α-tubulin-EGFP) and PH2B-mCherry-IRES-puro (RRID: Addgene_20972; H2B-mCherry) to monitor regular HEY cell division and cellular dynamics during PGCC endoreplication. Detailed information has been provided as described previously ^31^.

### 4.8 Wound healing assay

The daughter cells and clones were seeded into 6-well culture plates and allowed to reach approximately 95% confluence. A straight vertical scratch wound was then created by scraping with a sterile 200 µL pipette tip. Subsequently, the cells were incubated in medium containing 2% FBS to promote migration. Wound closure was monitored by capturing images at the specified time points, and the closure rate was quantified accordingly.

### 4.9 Transwell migration assay

For the migration assay, 1 × 10 cells suspended in 100 μL of serum-free medium were seeded into the upper chamber of a Transwell insert (8 μm pore size; Merck-Millipore, Darmstadt, Germany) and allowed to migrate for 18 hours. The lower chamber was filled with 500 μL of complete medium supplemented with 10% FBS to create a chemotactic gradient. Following migration, non-migrated cells remaining on the upper surface of the membrane were gently scraped off with a sterile cotton swab. Cells that had migrated through the membrane to the lower surface were fixed with 4% paraformaldehyde and stained with 0.05% crystal violet for 15 minutes. The average number of migrated cells per insert was calculated to quantify migration efficiency.

### 4.10 Immunofluorescence staining

The cells seeded on round glass coverslips were fixed with 1% (w/v) paraformaldehyde for 30 minutes, then incubated in blocking solution (3% BSA and 0.2% Triton X-100 in PBS) at room temperature for 1 hour. Primary antibodies were diluted in blocking solution (2% BSA in PBS) at a 1:100 dilution and applied to the cells, followed by an overnight incubation at 4°C. The next day, fluorophore-conjugated secondary antibodies (FITC- or PE-conjugated; diluted in 2% BSA in PBS; 1:200; Cell Signaling Technology, Danvers, MA) were added and incubated with the cells at room temperature for 30 minutes. Subsequently, the cells were stained with 4’,6-diamidino-2-phenylindole (DAPI, Beyotime) for 2 minutes at room temperature. Finally, the slides were mounted using fluorescent mounting medium (Agilent-DAKO, Santa Clara, CA) and visualized under a fluorescence microscope (Eclipse TE 2000-U; RRID: SCR_023161; Nikon, Tokyo, Japan).

### 4.11 SA-**β**-gal staining for senescence

For senescence assessment via SA-β-gal staining, cells were processed using a β-galactosidase staining kit (Beyotime, Shanghai, China). Briefly, cells were washed three times with phosphate-buffered saline (PBS) (5 minutes per wash), fixed with 1× fixative solution from the kit, and then rinsed again with PBS three times (3 minutes per wash). Staining was performed using the kit’s staining solution under dark conditions for 16 hours at 37°C. Finally, stained cells were observed and counted under an Axio Observer microscope (RRID: SCR_021351; Carl Zeiss, Oberkochen, Germany).

### 4.12 Spheroid formation assay

To assess the temporal dynamics of stemness in VCR-induced PGCCs (harvested at indicated times), we seeded 500 PGCCs/well (low-density) into 6-well ultra-low-attachment plates with stem cell medium (DMEM/F12 (BI) + 20 ng/mL bFGF (STEMCELL Technologies, Vancouver, Canada) + 20 ng/mL EGF (STEMCELL Technologies) + 1× B27 (STEMCELL Technologies) for 10/12 days. Regular HEY cells under identical conditions exhibited negligible spheroid formation (>200 μm) as a baseline control. Spheroids were observed and counted under an Axio Observer microscope. Single PGCC spheroidogenesis was evaluated by seeding 1–2 PGCCs/well into 96-well ultra-low-attachment plates. Spheroid formation was monitored after 16-day culture. For the spheroid formation capacity analysis of PGCCs-derived daughter cells, 500 cells/well or 3000 cells/well were cultured in ultra-low attachment 6-well plates for 14 days.

### 4.13 Trilineage differentiation assay

Pre-budding stage PGCCs (500 cells per well) and regular HEY cells (5,000 cells per well) were seeded onto ultra-low attachment 6-well plates. Cells were initially cultured in the aforementioned stem cell medium for 3 days to maintain their undifferentiated state and promote aggregation. Subsequently, the culture medium was replaced with STEMdiff™ Trilineage Media (STEMCELL Technologies), and differentiation was induced under lineage-specific conditions: endoderm lineages were cultured for 1, 3, 5, and 10 days supplemented with 200 ng/ml NOG (MCE) and 25 ng/ml Wnt3a (Beyotime). For the 10-day group, to enhance tissue differentiation, Corning® Matrigel® Matrix (Corning, NY) was further added to the medium on day 5 at a 1:1 ratio, along with 10 μM Y-27632, while the mesoderm and ectoderm lineage was cultured for 5 and 7 days, respectively. Spheroid formation and growth were monitored throughout the differentiation period. After the culture period, the spheroids were subjected to H&E staining.

### 4.14 Cell differentiation to neuronal lineage commitment

PGCCs (D10 induced with 1 μM VCR; 3000 cells/well in 6-well plates) with STEMdiff™ Forebrain Neuron Differentiation medium (Stemcell, Cambridge, MA) for 12 days and STEMdiff™ Forebrain Neuron Maturation medium (Stemcell) for 14 days following the manufacturer’s instructions, with with concurrent inhibition of BMP/SMAD signaling throughout the differentiation and maturation processes (0.5 μM of LDN-193189 (MCE) and 10 μM of SB431542 (MCE)).

### 4.15 Induction of adipogenesis in PGCCs

PGCCs (D10 induced with 1 μM VCR; 3000 cells/well in 6-well plates) were cultured with Adipogenic Differentiation A medium (Procell, Wuhan, China) for 2 days and Adipogenic Differentiation A medium (Procell) for 1 day, repeated three times following the manufacturer’ s instructions. After differentiation, cells were fixed in 4% formalin for 30 min and stained with Oil Red O. Oil Red O was extracted by isopropyl alcohol, and its optical density was monitored spectrophotometrically at 520 nm.

### 4.16 Animal experiments

All mouse experiments were approved by the Institutional Animal Care and Use Committee of SUSTech (Resolution number: SUSTech-JY202103028-A1). Twenty female BALB/c nude mice (6-8 weeks old) were obtained from Charles River (Beijing, China) and maintained in the Animal Center of SUSTech for one week before use. The daughter cells at the 27^th^ generation, derived from 1 μM VCR-induced PGCCs and regular HEY cells, were subcutaneously injected into the right flank area of each mouse for 33 days (5 × 10^6^ cells/mouse; 10 mice/group). Tumor volumes were measured (0.5 × length × width^2^) every 5 to 7 days after palpable tumors appeared. On day 33, mice were euthanized and tumors were surgically isolated, photographed, and weighed.

### 4.17 RNA sequencing

The RNA sequencing experiments were conducted by LC-Bio Co. Ltd. (Hangzhou, China). Briefly, total RNA was extracted from VCR-induced PGCCs at D10 and regular HEY cells in biological triplicate using TRIzol reagent (Invitrogen, Carlsbad, CA, USA) and then subjected to library construction. The detailed protocols for subsequent sequencing and data analysis were performed as previously reported ^32^. The dataset has been submitted to the NCBI GenBank under BioProject ID PRJNA1204050, which has been released on December 31, 2025.

### 4.18 qRT-PCR

To validate the RNA sequencing results, quantitative real-time PCR (qRT-PCR) was performed for the differential mRNAs of SASP-associated cytokines. GAPDH was used as the endogenous reference gene. Total RNA for qRT-PCR was collected as described above. cDNA was synthesized from total RNA using the M-MLV RTase cDNA Synthesis kit (Takara, Dalian, China) according to the manufacturer’s instructions. Quantitative real-time PCR was performed on a qTOWER^3^ (RRID: SCR_027122; Analytik Jena, Jena, Germany) using SYBR Premix Ex Taq (Takara, Dalian, China) according to the manufacturer’s protocol. The primer sequences are listed in the **Supplementary Table 2** (**Table S2**).

### 4.19 Transfection of siRNAs

SiRNA transfection was performed as previously described ^33^. The sequences of siRNA oligonucleotides are listed in the **Supplementary Table 3** (**Table S3**).

### 4.20 Selection and subculture of single PGCC-derived daughter cell clones

His2B-labeled HEY cells were treated with VCR for 2 days, followed by a 10-day recovery period. The VCR-induced PGCCs were then subcultured into 96-well plates at a density of 0.5 cells per well. After 2 days of culture, wells containing only a single attached PGCC were identified and marked. Colonies were monitored every 3 days, and wells with budding PGCCs were recorded. Two days later, single PGCC-derived daughter cell clones were subcultured into 24-well plates for subsequent experiments.

### 4.21 Human XL Cytokine Array

Supernatants from the PGCCs at D10 and regular HEY cells were analyzed using the Human XL Cytokine Array Kit (ARY022; R&D Systems, Minneapolis, MN) according to the manufacturer’s instructions.

### 4.22 Enzyme-linked immunosorbent assay (ELISA)

IL1α (Catalog no. KE00268; Proteintech, Wuhan, China), IL1β (Catalog no. orb50042; Biorbyt, Cambridge, UK), IL6 (Catalog no. SEKH-0013; Solarbio, Beijing, China), IL8 (Catalog no. SEKH-0016; Solarbio, Beijing, China) and CXCL1 (Catalog no. KE00133; Proteintech, Wuhan, China) levels in supernatants of PGCCs (D16), PGCCs with budding (D36) and regular HEY cells were detected by ELISA kits according to the instructions of the manufacturer.

### 4.23 Immunocytochemical staining

PGCCs (D16), budding PGCCs (D36), and regular HEY cells cultured in 12-well plates were fixed with 1% paraformaldehyde, followed by two washes with 1× PBS. Subsequently, cells were permeabilized with 0.2% Triton X-100 in PBS and blocked with 3% BSA in PBS for 30 minutes at room temperature. Primary antibodies were then applied, and the samples were incubated at 4°C overnight. After three washes with 1× PBS, cells were processed using a DAB Detection System (MXB, Fuzhou, China) according to the manufacturer’s instructions. Nuclei were counterstained with hematoxylin (MXB) for 1 minute. Finally, stained cells were observed and counted under an Axio Observer microscope.

### 4.24 Chemosensitivity assay

Cells were seeded into 96-well plates at a density of 5 × 10^3^ cells per well and allowed to adhere overnight. Subsequently, cells were treated with varying concentrations of VCR for an additional 48 hours. Cell viability was assessed using a CCK-8 assay kit (Beyotime) according to the manufacturer’s instructions. Absorbance values were normalized to those of the DMSO-treated control group, and IC_50_ values were calculated using GraphPad Prism 8.0.1 (RRID: SCR_002798; GraphPad Software, La Jolla, CA).

### 4.25 Statistical analyses

All experimental data were analyzed using GraphPad Prism 8.0.1. All statistical analyses employed two-tailed Student’s *t*-tests unless otherwise specified in the figure legends. Values of *p* < 0.05 were considered statistically significant.

## Acknowledgments

This work was supported by the Institutional incentive salary and Briding Fund, a fund from OvaCure, and National Natural Science Foundation of China (32170568), National Natural Science Foundation of China for Youth (82002997), General Project of Basic Research in Shenzhen (JCYJ20190809161215057), Medical Scientific Research Foundation of Guangdong Province of China (A2022131) and Shenzhen Medical Research Fund (B2402039).

## Author contributions

JL and ZZ conceived and designed the experiments. JD participated in conceptual discussions and experiment design. ZZ performed most of the experiments. XL performed time-lapse image acquisition. XT conducted qPCR and a subset of Western blot analyses. LD assisted with specific steps in cell sample preparation. ZZ drafted the manuscript. JL, ZZ and JD revised the manuscript. JL and ZZ finalized the manuscript.

