## Supplemental data legends for "Therapeutic Stress-induced Activation of PGCC Life Cycle Drives the Resistance Acquisition and Structured Tissue Differentiation"

**Supplemental Figure 1. VCR-induced PGCC formation and budding dynamics in HEY cells across varying concentrations. (A and B)** The HEY cells were treated with different concentrations of VCR (A: 0.5 μM, B: 4 μM) for 2 days, then allowed to recover for PGCC formation and budding into daughter cells. For each figure, the upper panel indicates the morphology of PGCCs at different time points. Upper lane: phase contrast images; middle lane: histone 2B-GFP labeled to show the nuclei; lower lane: merged images. The lower panel shows the cell cycle distribution and percentage of cells with a DNA content greater than 4N in HEY cells treated with the corresponding concentration of VCR for 2 days and then allowed to recover in drug-free culture medium for the indicated days. Scale bars: 200 μm. **(C and D)** Crystal violet staining to show the budding daughter cell clones from different concentrations of VCR-induced PGCCs (upper left: 0.5 μM; upper middle: 1 μM; upper right: 4 μM). Scale bars: 10 mm. C: representative images; D: the number of budding daughter cell clones for different concentrations of VCR induced PGCCs (3000 PGCCs per well). The experiment was performed in triplicate. Data are presented as mean ± SEM. * *p* < 0.05, ** *p* < 0.01, *** *p* < 0.001.

**Supplemental Figure 2. VCR-induced PGCC formation and budding dynamics in SKOV3 cells at varying concentrations. (A-C)** SKOV3 cells were exposed to VCR at indicated concentrations (A: 0.5 μM, B: 1 μM, C: 4 μM) for 2 days, followed by a recovery period to allow PGCC formation and budding into daughter cells. For each subfigure, the upper panel illustrates the morphological progression of SKOV3 cells during PGCC formation and budding. The upper lane displays phase-contrast microscopy images; the middle lane shows Hoechst 33342-stained nuclei; the lower lane presents merged images. Scale bars, 200 μm. The lower panel quantifies the cell cycle distribution and the proportion of cells with a DNA content greater than 4N in SKOV3 cells following VCR treatment (as indicated) for 48 hours, with recovery periods as specified. **(D)** Crystal violet staining visualizes budding progeny clones derived from VCR-induced PGCCs at different concentrations (upper left: 0.5 μM; upper middle: 1 μM; upper right: 4 μM). The lower panel quantifies the number of budding progeny clones per well (1500 PGCCs seeded per well) across VCR concentrations. The experiment was performed in triplicate. Data are presented as mean ± SEM. * *p* < 0.05, ** *p* < 0.01, *** *p* < 0.001.

**Supplemental Figure 3. Microscopic visualization confirmed endoreplication and polyploidization in HEY cells exposed to 1 μM VCR. (A, B)** Time-lapse imaging acquired through visual monitoring and manual operational control to monitor endoreplication in VCR-induced PGCCs derived from HEY cells expressing the Fucci cell cycle-visualization system. Upper panel: Representative time-lapse frames showing a single PGCC in the G1/S phase with progressive DNA content increase. Lower panel: Representative time-lapse frames depicting the endomitotic process of a single cell, progressing through the G1/S→S/G2→G1/S phases. Scale bars: 200 μm. **(C, D)** Quantification of single PGCC nuclear volume over the indicated time points in (A) and (B), respectively.

**Supplemental Figure 4. Comparative analysis of replication dynamics in normal vs. PGCCs.** Stable HEY cells expressing H2B-mCherry (histone 2B) and α-tubulin-GFP (microtubules) were imaged to visualize replication machinery. Left panel: Normal mitosis. Right panel PGCC endomitosis.

**Supplemental Figure 5. Daughter cells derived from VCR-induced PGCCs progressively acquire enhanced proliferative capacity and exhibit enhanced migratory capacity compared to regular HEY cells.** **(A)** Proliferation rates of 3^rd^- (left) and 27^th^- (right) passage daughter cells versus regular HEY cells were quantified using SRB assays. Data are presented as mean ± SEM. * *p* < 0.05. **(B)** Drug sensitivity profiling of VCR in regular HEY cells and 27^th^-passage daughter cells derived from different concentrations of VCR-induced PGCCs. Data presented as mean ± SEM. * *p* < 0.05, ** *p* < 0.01. **(C)** Representative images illustrate the migratory behavior of 27^th^ generation daughter cells derived from HEY cells, assessed via a scratch wound healing assay following induction with varying concentrations of VCR. **(D)** IF assay for EMT markers Pan-CK and FN1 in 27^th^-passage daughter cells derived from different concentrations of VCR-induced PGCCs in HEY cells. Scale bars: 100 μm.

**Supplemental Figure 6. Clonal isolation and motility profiling of single PGCC-derived daughter cell populations. (A)** Schematic illustration of the single PGCC-derived daughter cell clone isolation protocol. **(B)** Morphological analysis of a single PGCC at D10 (plated in the 96-well dish) derived from HEY cells induced by different VCR concentrations (0.5, 1, or 4 μM) and their progeny clones. Upper panels: Representative single PGCCs; Lower panels: Budding daughter cell clones. Left: Progeny clone 1 derived from a 0.5 μM VCR-induced single PGCC; Middle: 1 μM; Right: 4 μM. Scale bars: 200 μm. **(C-D)** Representative images of wound healing assay to evaluate the migratory capacity of progeny clones from different VCR-induced single PGCCs. Scale bars: 200 μm.

**Supplemental Figure 7. PGCCs induced by higher concentrations of VCR exhibit enhanced stemness.** **(A)** PGCCs induced by 0.5, 1, and 4 μM VCR were plated at D16 into ultra-low attachment 6-well plates at a density of 500 cells/well. Spheroid formation was observed after 12 days of culture in stem cell medium. **(B)** PGCCs induced by 0.5, 1, and 4 μM VCR at D16 were seeded into ultra-low attachment 96-well plates at a density of 1-2 cells/well. Spheroid formation was monitored after 16 days. **(C)** PGCCs induced with 0.5 μM VCR and harvested at D18, those induced with 1 μM VCR and harvested at D22, and those induced with 4 μM VCR and harvested at D26 were separately seeded into ultra-low attachment 6-well plates at a density of 500 cells per well. **(D)** PGCCs induced with 0.5, 1, or 4 μM VCR and harvested at D18, D22, or D26, respectively, were separately seeded into ultra-low attachment 96-well plates at 1-2 cells/well. **(E)** Pre-budding PGCCs (from 4 μM VCR) differentiated in endodermal medium yielded a mature epithelial chambered structure (this result was observed only once in an experiment).

**Supplemental Figure 8. Schematic illustrates that PGCCs exhibit dynamic changes in proliferation, EMT, and stemness properties.** Stress-induced cell cycle arrest in maternal PGCCs persists in early daughter cells, with their mitotic activity surpassing that of regular HEY cells after prolonged passaging. PGCCs maintain consistently enhanced EMT properties compared to HEY cells, with daughter cells further amplifying these EMT phenotypes. Stemness progresses biphasically: weak pluripotency initially, intensifying to blastomere-like functionality pre-budding, then attenuating post-budding.

**Supplemental Figure 9. Arrangement of 1d, 3d, 5d, and 10d spheroids by size and differentiation degree reveals comprehensive differentiation process.** Scale bars: 200 μm.

**Supplemental Figure 10. Central clearances (marked by arrows) are observed in the lumen formation of human tumors. (A)** High-grade serous carcinoma. **(B)** Endometrial hyperplasia. **(C and D)** Endometrioid adenocarcinoma. Scale bars: 100 μm.

**Supplemental Figure 11. VCR-induced PGCCs show astrocyte-like differentiation (GFAP⁺/TUBB3⁻).** PGCCs derived from VCR-treated HEY cells (D10) were subjected to culture in neuronal differentiation medium. **(A)** Immunofluorescence analyses of GFAP and TUBB3 expression in regular HEY cells. **(B)** Immunofluorescence analysis of GFAP and TUBB3 expression in differentiated PGCCs. Scale bars: 100 μm.

**Supplemental Figure 12. VCR-induced PGCCs exhibit SASP in SKOV3 cells. (A-B)** Detection of SASP-associated factors IL1α, IL1β, IL6, CXCL1, IL8/CXCL8, CSF3 and PTX3 in PGCCs at D10 and regular cells via qRT-PCR. Data are presented as mean ± SEM. * *p* < 0.05, ** *p* < 0.01. A: HEY cells; B: SKOV3 cells. **(C)** ELISA assay to determine the protein levels in the supernatants of PGCCs at D16 and D36 and regular HEY cells. Data are presented as mean ± SEM. ** *p* < 0.01; *** *p* < 0.001.

**Supplemental Figure 13. Validation of IL1**α**, IL1**β**, IL6, IL8, and CXCL1 Knockdown Efficiency via qRT-PCR.** Quantitative analysis of relative mRNA expression levels of IL1α, IL1β, IL6, IL8, and CXCL1 in HEY cells following RNA interference (RNAi). Data are presented as mean ± SEM. * *p* < 0.05, ** *p* < 0.01. ** *p* < 0.01.

**Supplemental Figure 14. Blocking IL1R1, IL-6R, and CXCR1/CXCR2 suppressed the PGCC formation and budding capacity in HEY and SKOV3 cells.** **(A-B)** IL1R1, IL-6R, and CXCR1/CXCR2 were antagonized with AF12198 (10 nM), Tocilizumab (1 μM), and Navarixin (1 μM), respectively. Then, 1 μM of VCR was added for 2 days and allowed to recover in a drug-free medium to induce the formation of PGCCs. Morphological analysis was performed at D0, D16 and D24. A: HEY cells; B: SKOV3 cells. Scale bars: 200 μm. **(C)** Quantification of PGCC numbers per well (D16) across each groups. Data are presented as mean ± SEM. * *p* < 0.05, ** *p* < 0.01. ** *p* < 0.001. **(D)** VCR-induced PGCCs (3000 per well) at D10 from HEY with IL1R1, IL-6R, and CXCR1/CXCR2 blocking were subcultured into 12-well plates and maintained for 30 days and the clone numbers were quantified. Data are presented as mean ± SEM. * *p* < 0.05. **(E)** IL1R1, IL-6R, and CXCR1/CXCR2 were antagonized with AF12198 (10 nM), Tocilizumab (1 μM), and Navarixin (1 μM) in HEY cells, respectively, then 1 μM of VCR was added for 2 days and allowed for recovery to induce the formation of PGCCs. Ten days later, these PGCCs were cultured in neuron differentiation and maturation medium following the aforementioned protocol. Subsequently, cell morphology was examined under bright-field microscopy.

**Supplementary Table 1. Antibodies used for western blot (WB), immunofluorescence (IF), and immunocytochemistry (ICC) assays.**

**Supplementary Table 2. qRT-PCR primers.**

**Supplementary Table 3: The sequences of small interfering RNA.**

**Supplemental movie 1. Original time-lapse time-lapse videos for Fig. 1C.**

**Supplemental movie 2. Original time-lapse videos for the left panel of Fig. S4.**

**Supplemental movie 3. Original time-lapse videos for the right panel of Fig. S4.**
