## Supplementary figures and images for "Therapeutic Stress-induced Activation of PGCC Life Cycle Drives the Resistance Acquisition and Structured Tissue Differentiation"

### Figure S1

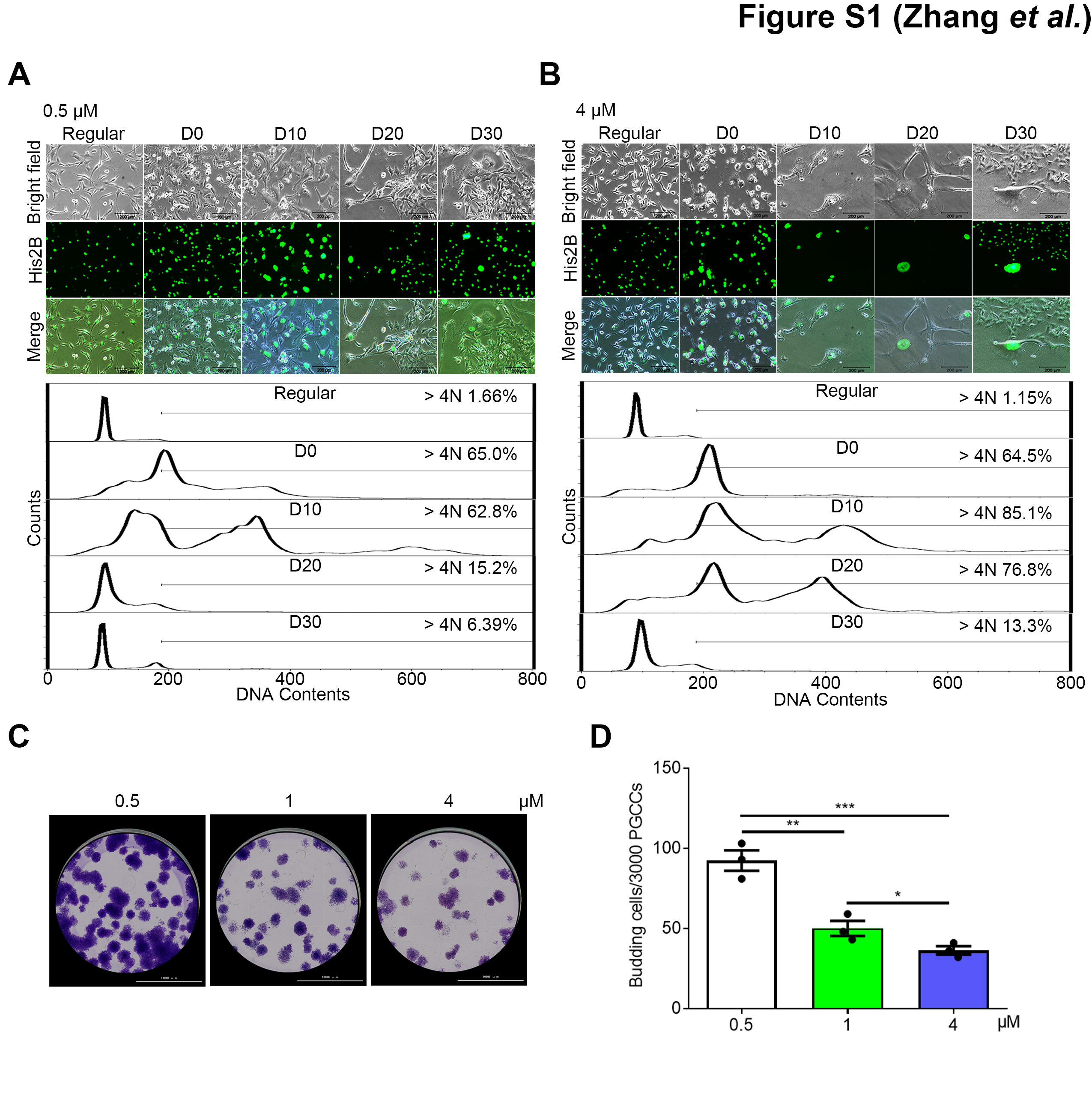

### Figure S2

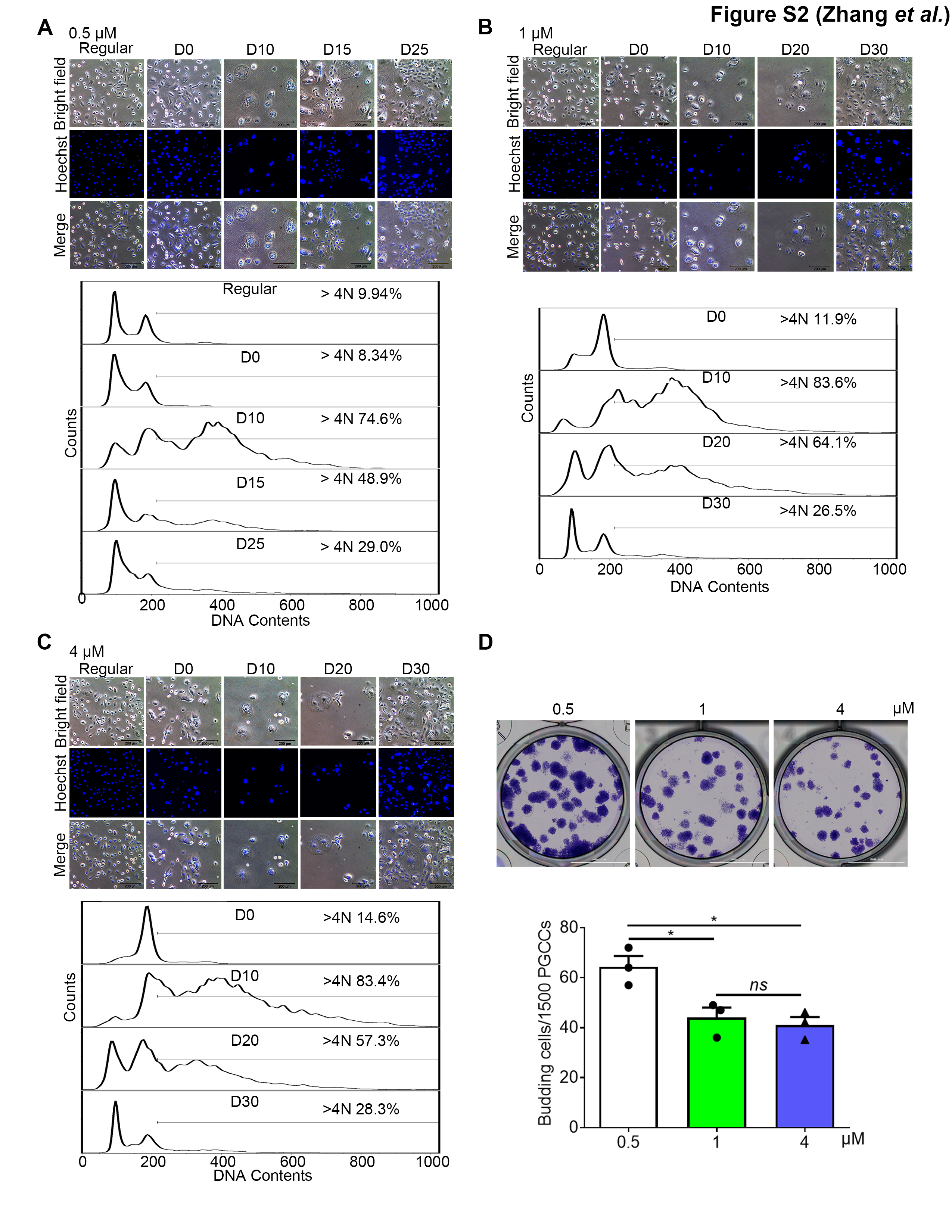

### Figure S3

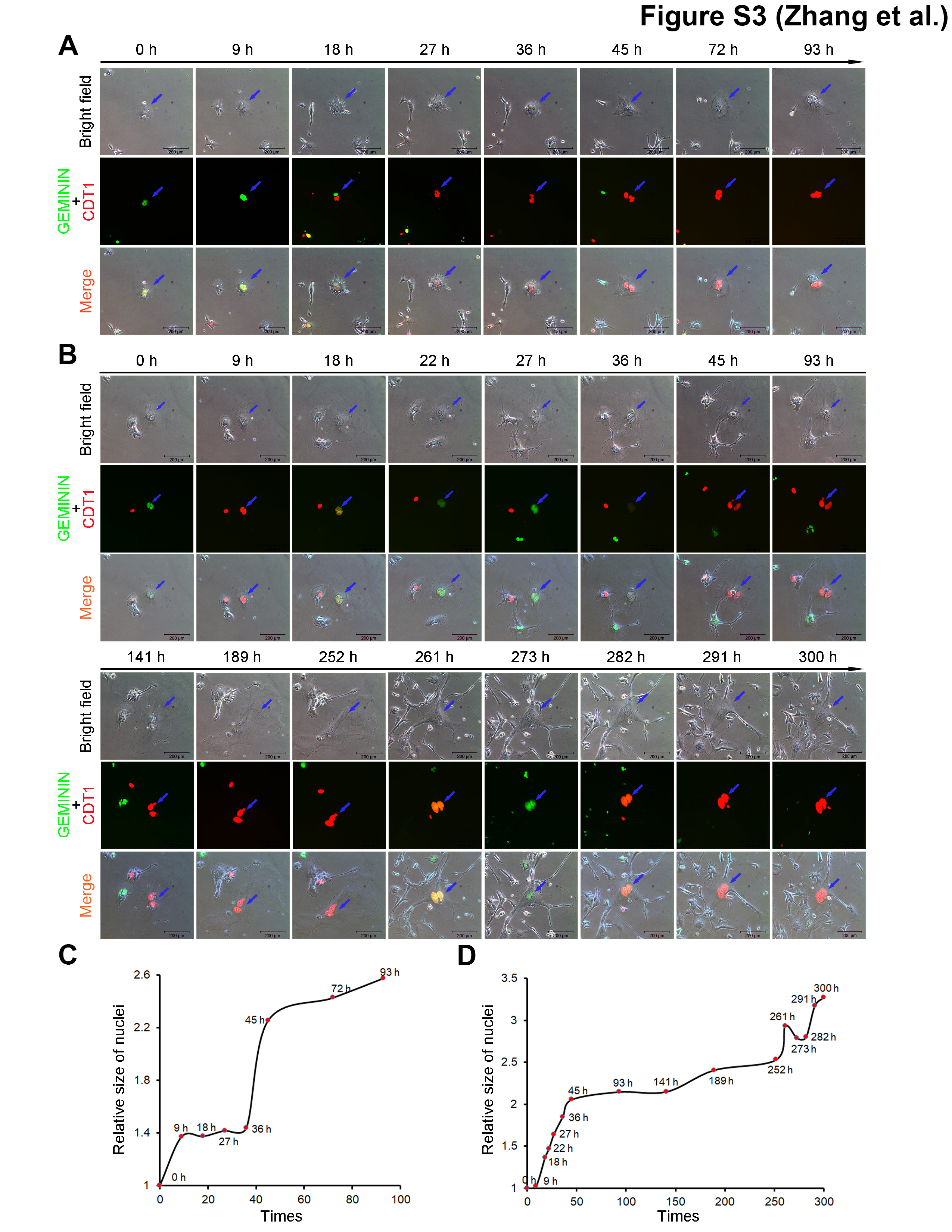

### Figure S4

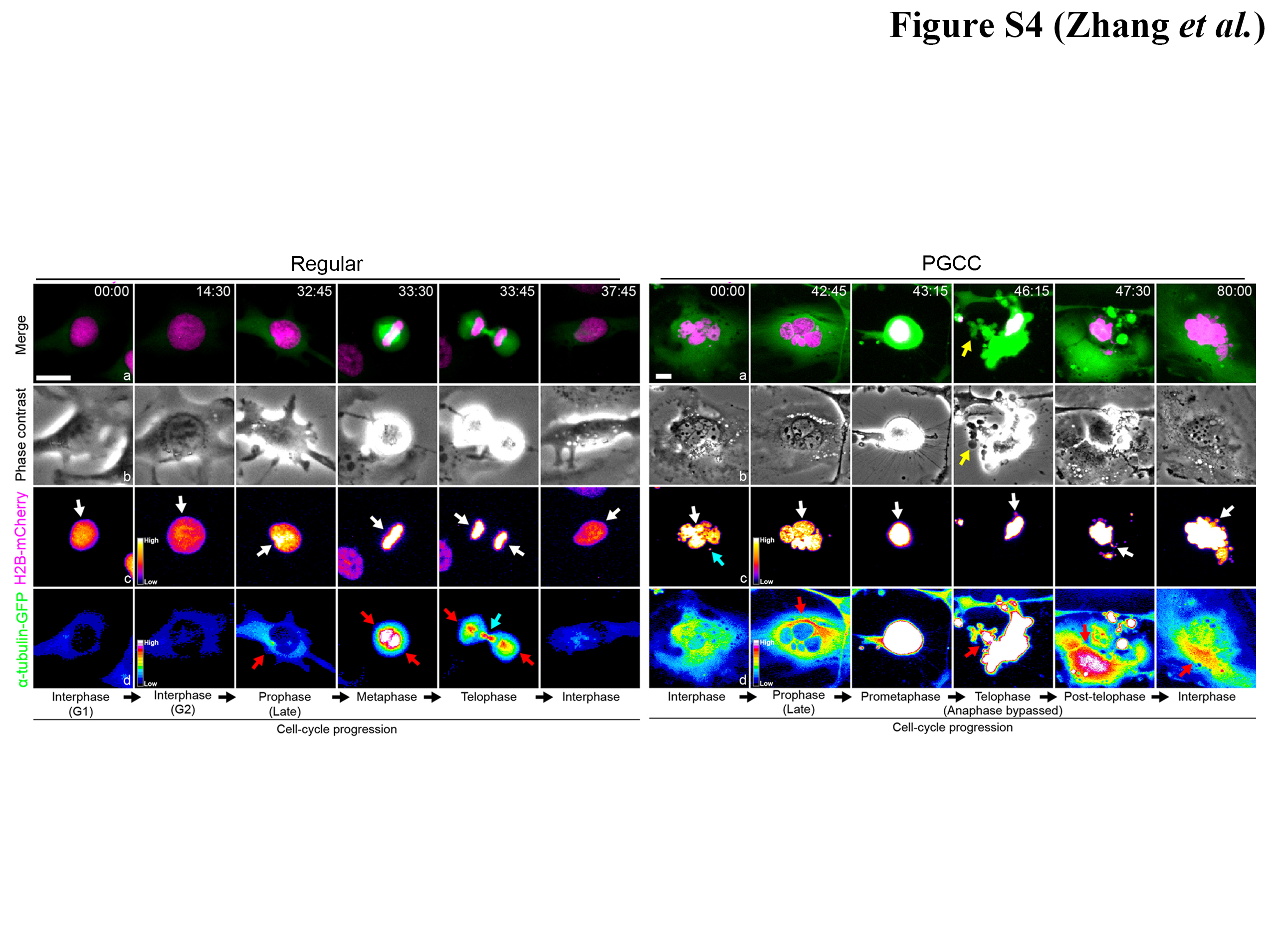

### Figure S5

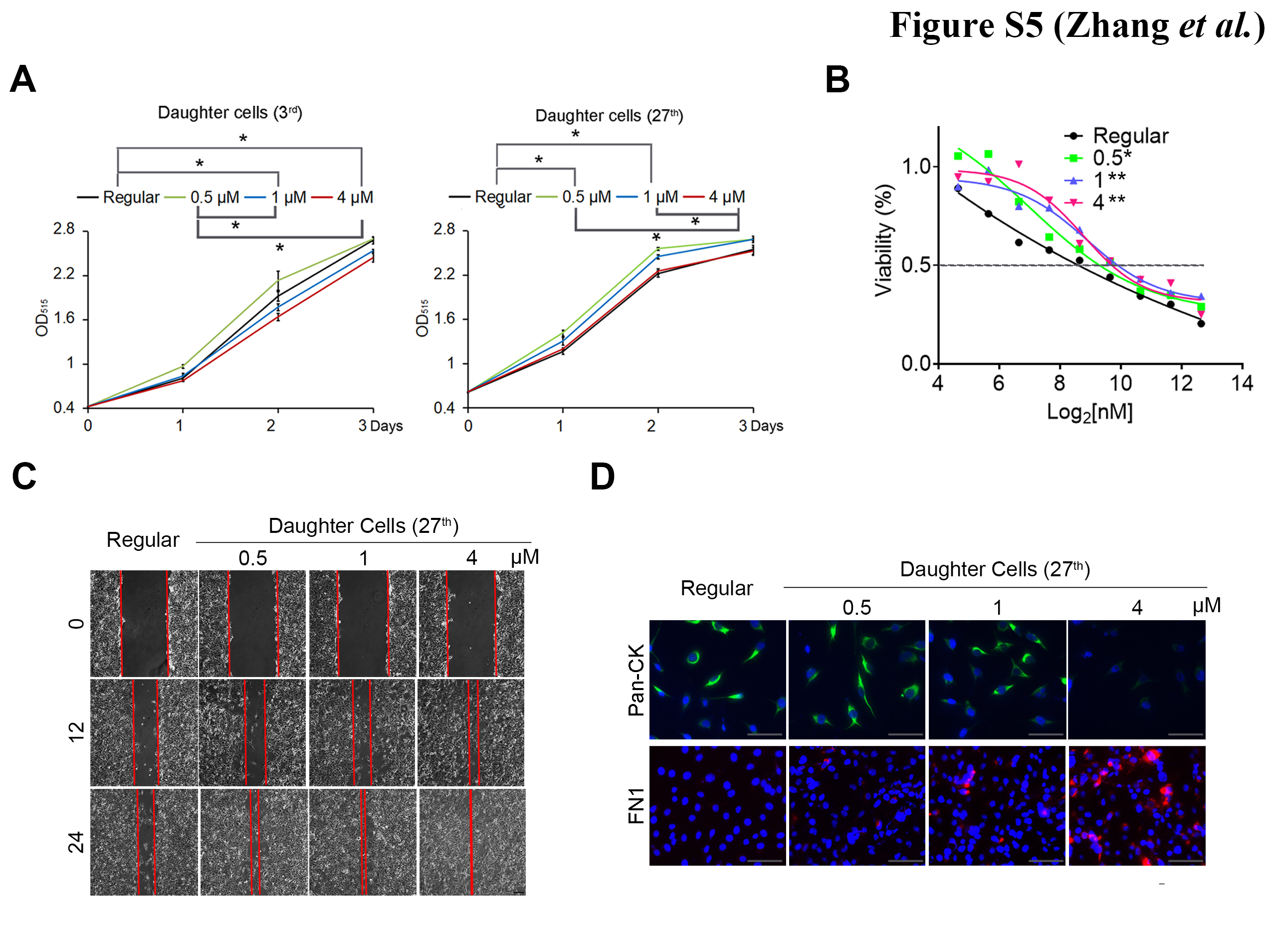

### Figure S6

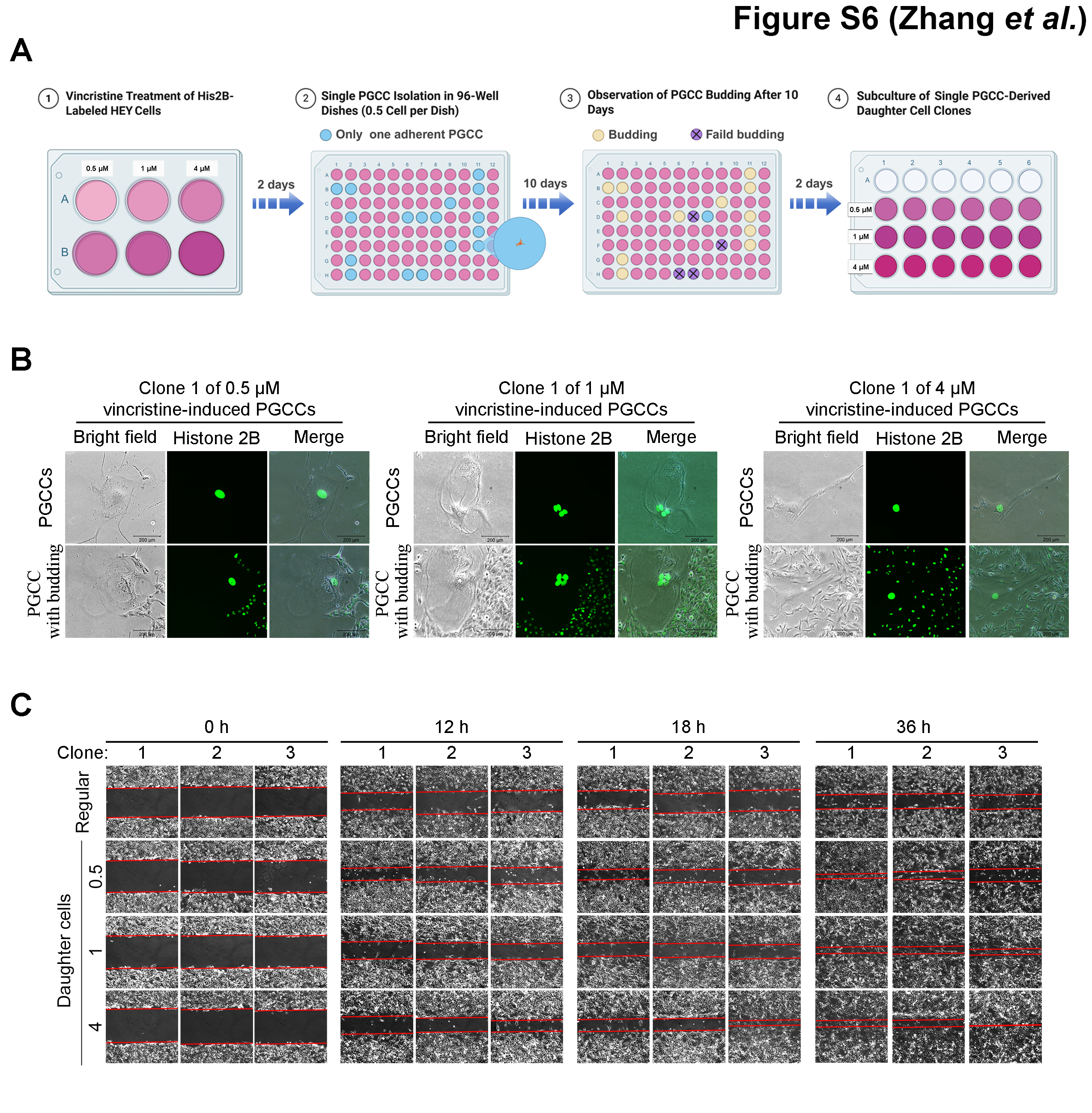

### Figure S7

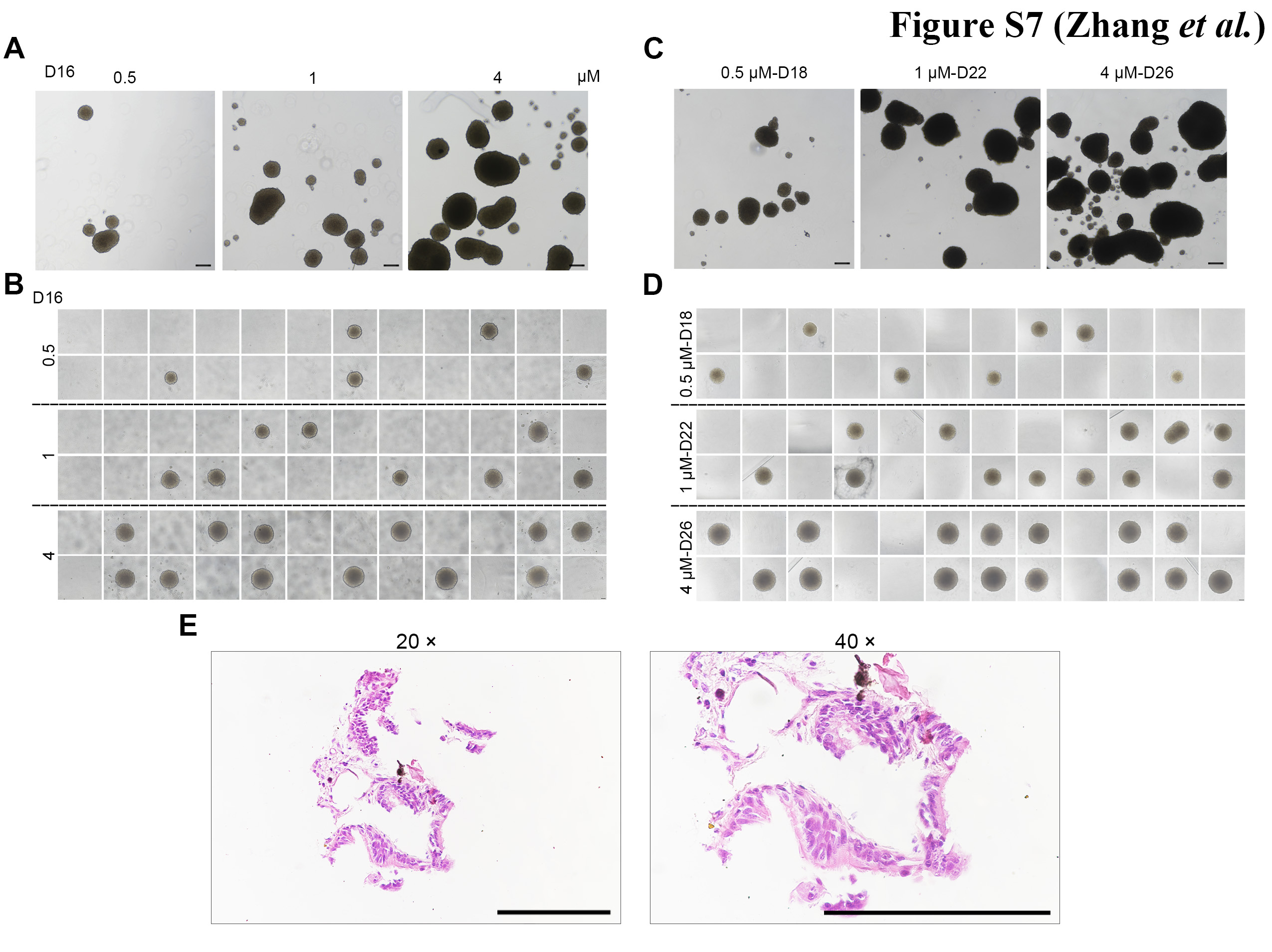

### Figure S8

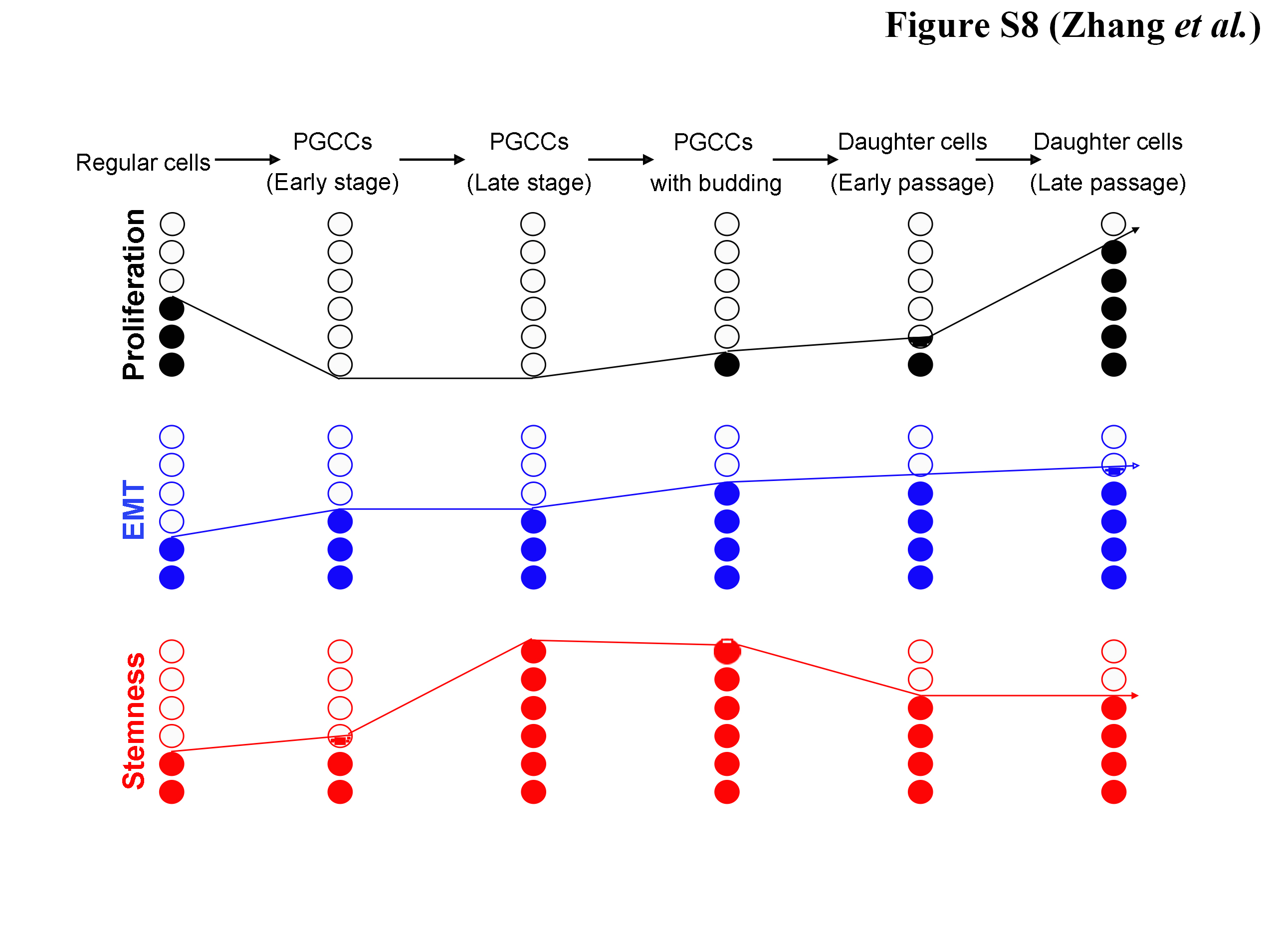

### Figure S9

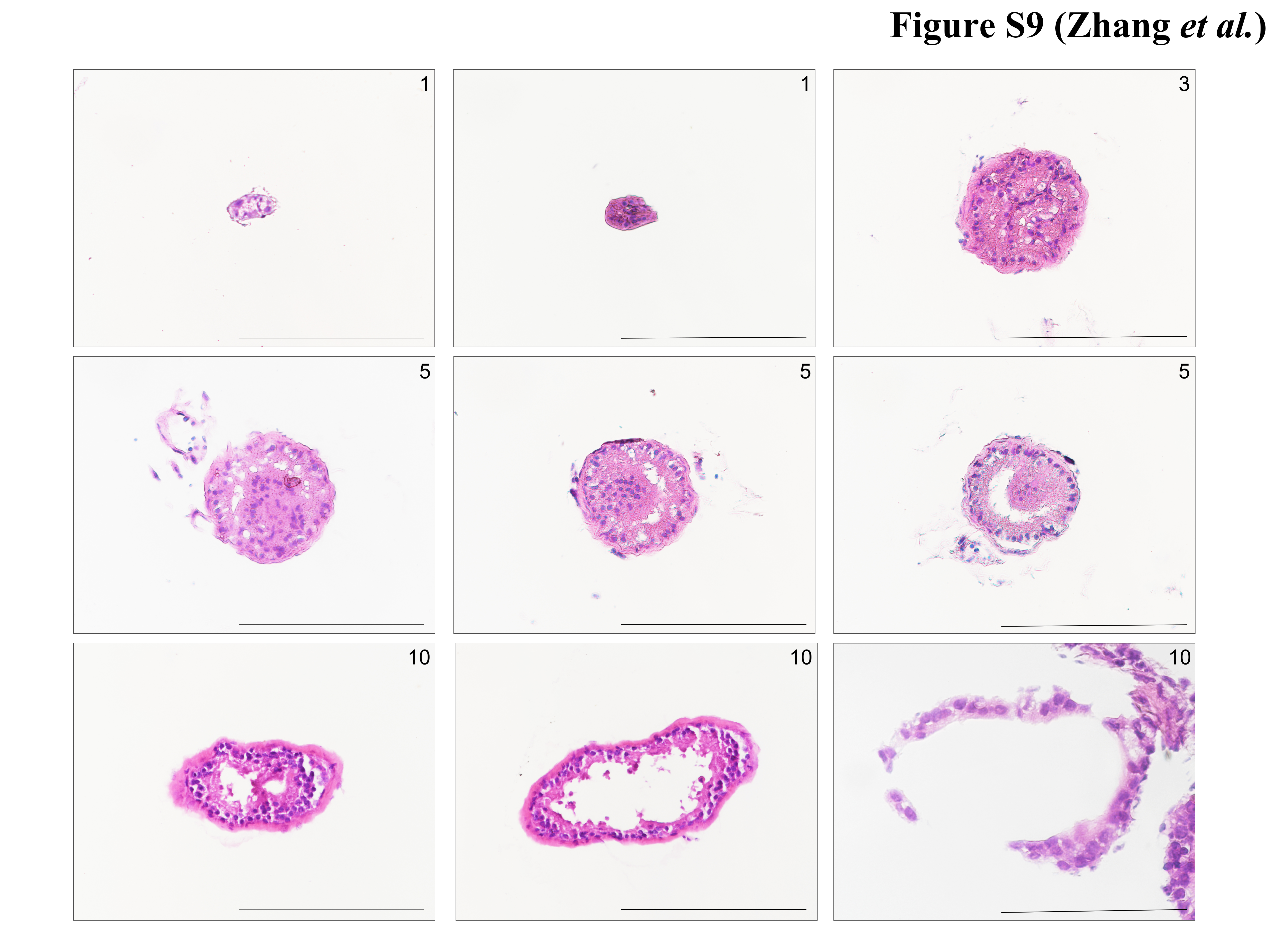

### Figure S10

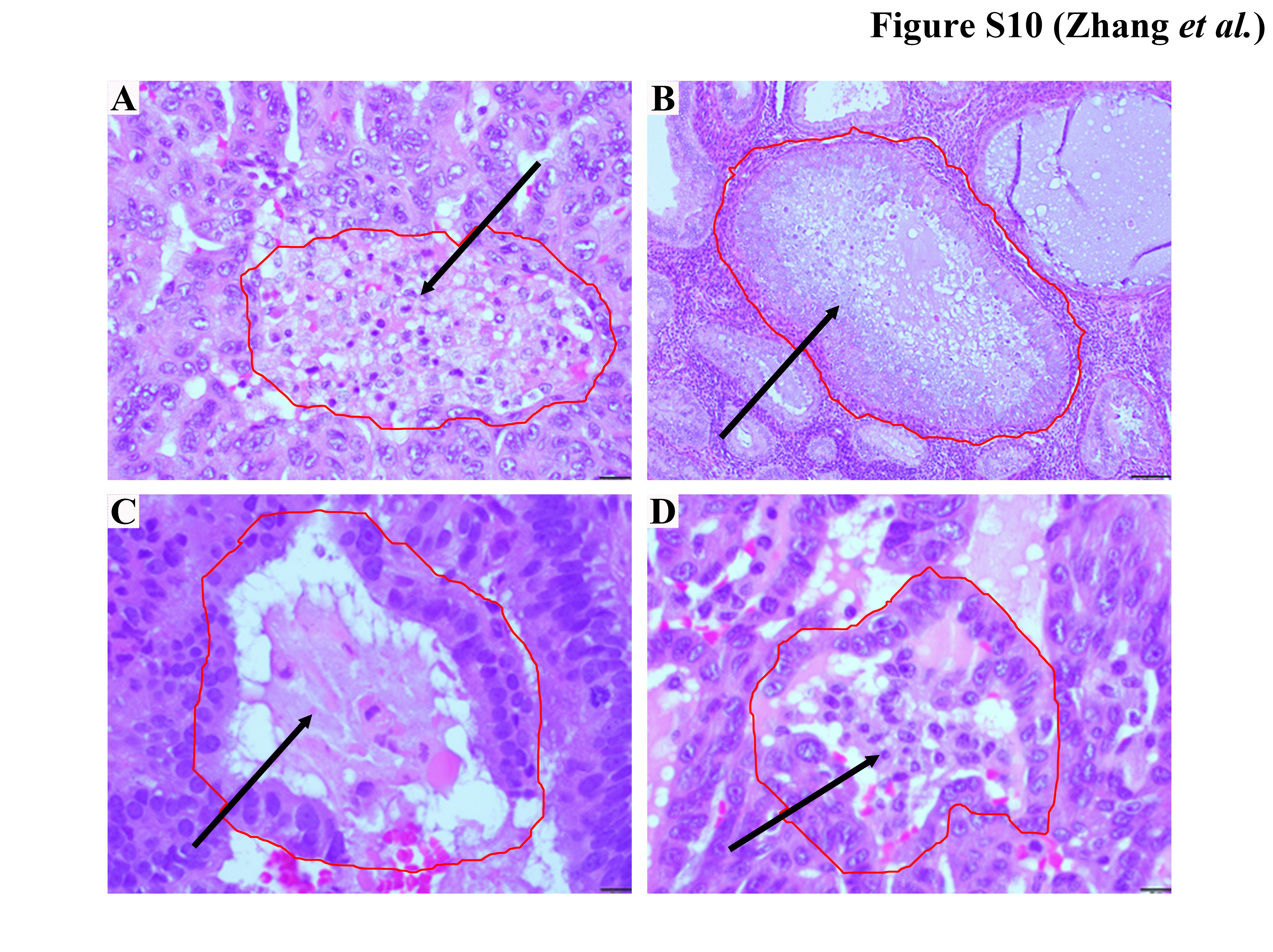

### Figure S11

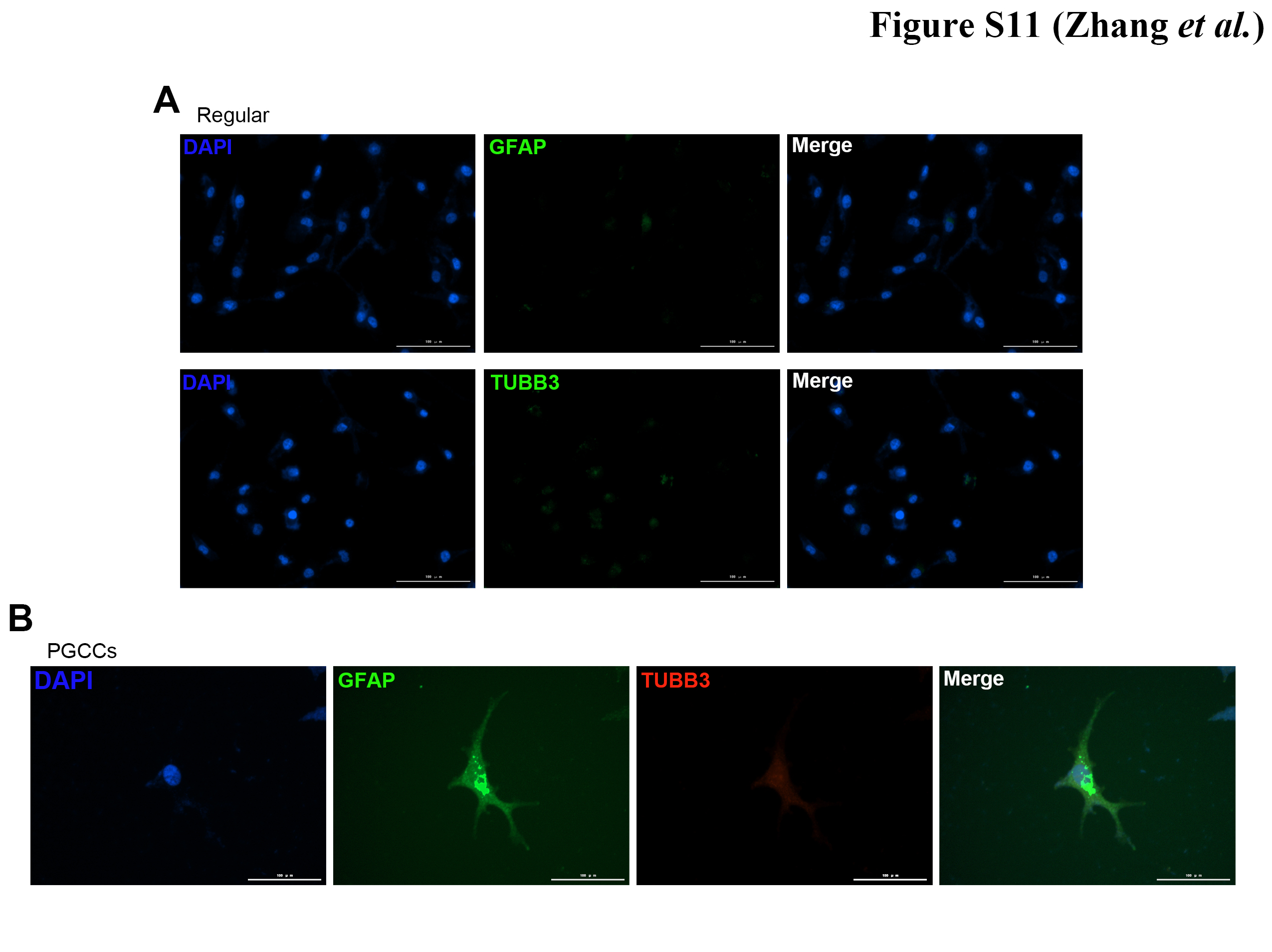

### Figure S12

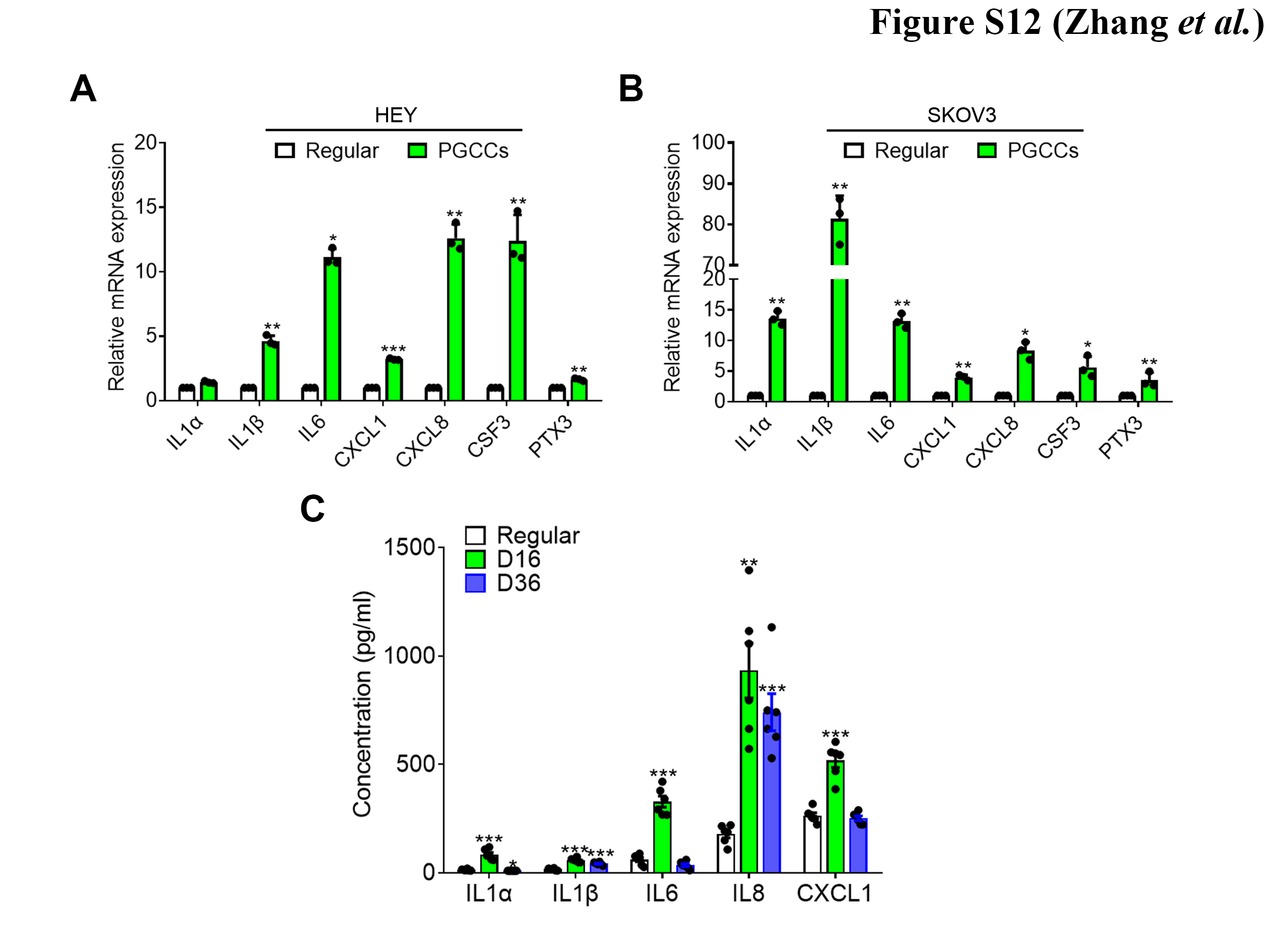

### Figure S13

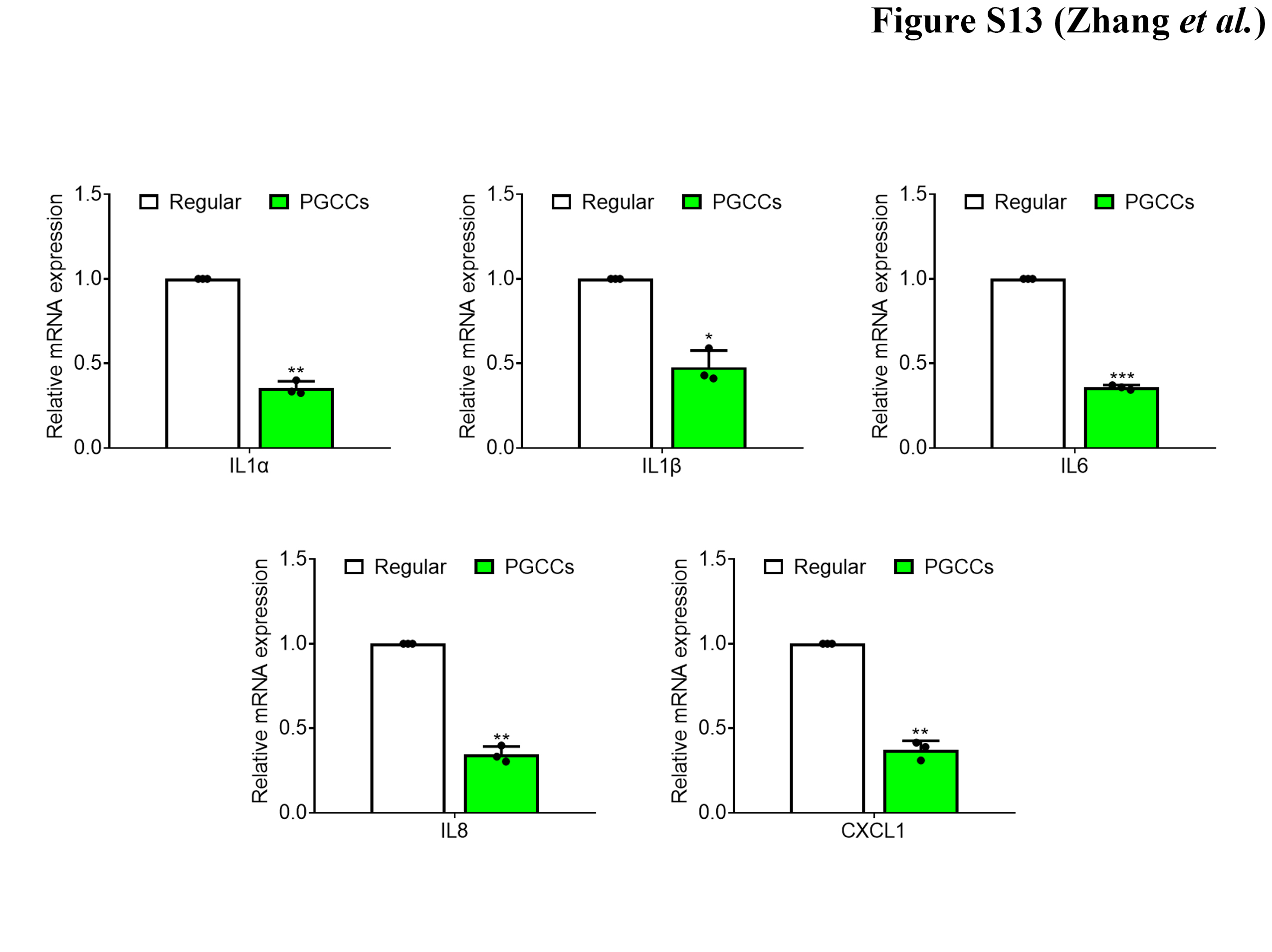

### Figure S14

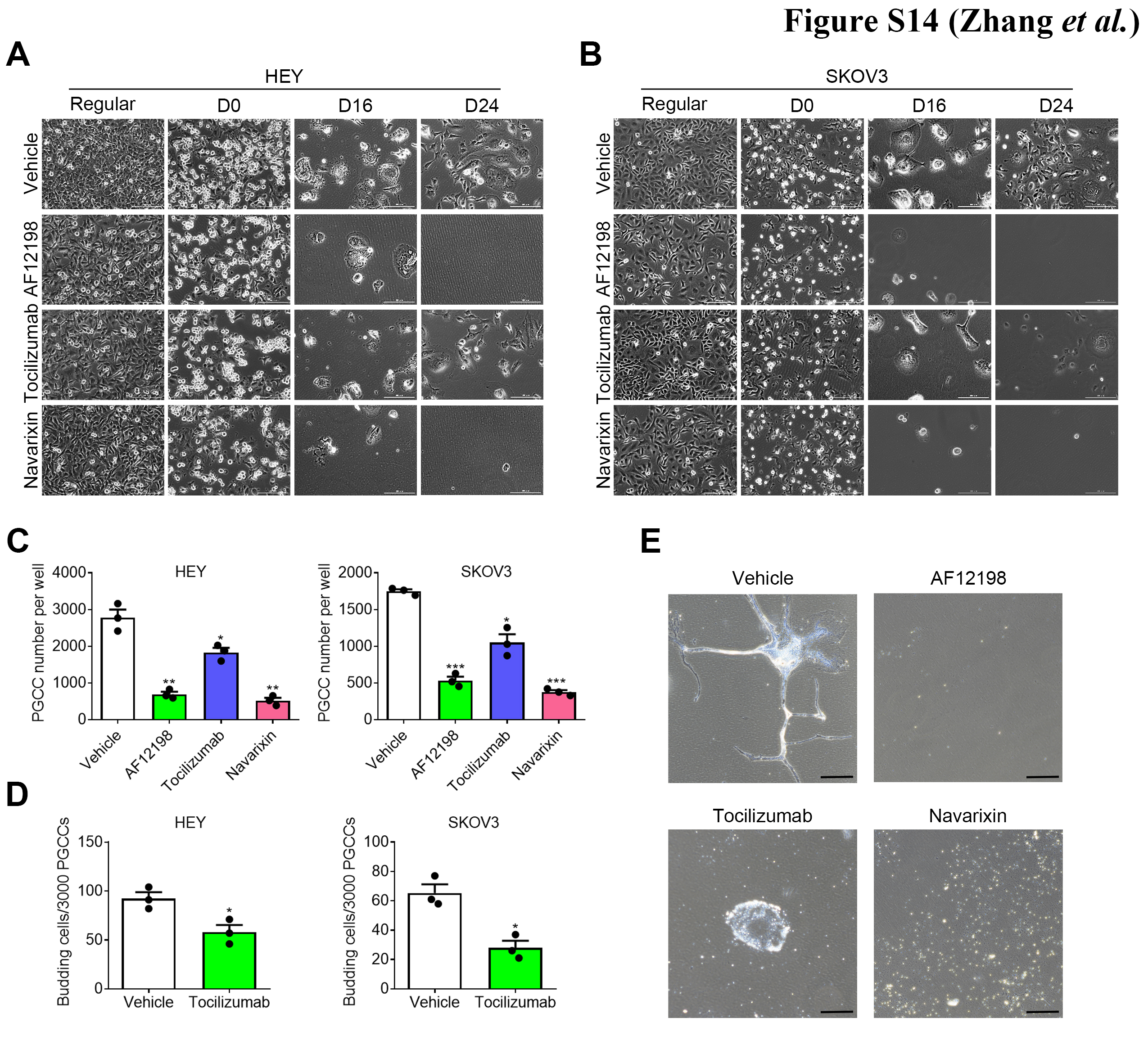
