## Supplemental table 1 for "Therapeutic Stress-induced Activation of PGCC Life Cycle Drives the Resistance Acquisition and Structured Tissue Differentiation"

| **Antibodies** | **Company** | **Catalog No.** | **RRID** | **Dilution** |
| --- | --- | --- | --- | --- |
| p16 | Cell Signaling | 80772 | AB_2799960 | WB: 1:1000 |
| Rb | Cell Signaling | 9313 | AB_1904119 | WB: 1:2000 |
| p-Rb | Cell Signaling | 9301 | AB_330013 | WB: 1:1000 |
| IL1α | Proteintech | 16765-1-AP | AB_10641044 | WB: 1:1000; ICC: 1:200 |
| IL1β | Cell Signaling | 12703 | AB_3083638 | WB: 1:1000; ICC: 1:200 |
| IL6 | Proteintech | 21865-1-AP | AB_11142677 | WB: 1:1000; ICC: 1:200 |
| IL8 | Proteintech | 27095-1-AP | AB_11142677 | WB: 1:1000; ICC: 1:200 |
| Cyclin A1 | Sigma | SAB4503500 | AB_10751191 | WB: 1:1000 |
| Cyclin B1 | Cell Signaling | 4138 | AB_2072132 | WB: 1:1000 |
| Cyclin D1 | Cell Signaling | 2978 | AB_2259616 | WB: 1:1000 |
| CXCL1 | Cell Signaling | 24376 | AB_3075424 | WB: 1:1000; ICC: 1:200 |
| Pan-CK | Proteintech | 26411-1-AP | AB_2919645 | WB: 1:1000;  IF: 1:200 |
| SNAIL | Cell Signaling | 3879 | AB_2255011 | WB: 1:1000 |
| CDH2 | Cell Signaling | 13116 | AB_2687616 | WB: 1:1000 |
| FN1 | Cell Signaling | 26836 | AB_2924220 | WB: 1:1000;  IF: 1:200 |
| GFAP | Cell Signaling | 80788 | AB_2799963 | WB: 1:1000;  IF: 1:200 |
| TUBB3 | Proteintech | 66375-1-1g | AB_2814998 | IF: 1:200 |
| EpCAM | Cell Signaling | 36746 | AB_2799105 | IF: 1:200 |
| β-ACTIN | Sigma | A2228 | AB_476697 | 1:3000 |

**Supplementary table 1. Antibodies used for western blot (WB), immunofluorescence (IF), and immunocytochemistry (ICC) assays.**
