## Supplemental table 2 for "Therapeutic Stress-induced Activation of PGCC Life Cycle Drives the Resistance Acquisition and Structured Tissue Differentiation"

| **Gene** | **Forward primer** | **Reverse primer** |
| --- | --- | --- |
| IL1α | AGATGCCTGAGATACCCAAAACC | CCAAGCACACCCAGTAGTCT |
| IL1β | ATGATGGCTTATTACAGTGGCAA | GTCGGAGATTCGTAGCTGGA |
| IL6 | ACTCACCTCTTCAGAACGAATTG | CCATCTTTGGAAGGTTCAGGTTG |
| IL8/CXCL8 | TGTGAAGGTGCAGTTTTGCCAAGG | GTTGGCGCAGTGTGGTCCACTC |
| CXCL1 | CTTGCCTCAATCCTGCATC | CCTTCTGGTCAGTTGGATTTG |
| CSF3 | ATAGCGGCCTTTTCCTCTACC | GCCATTCCCAGTTCTTCCAT |
| PTX3 | AGGCTTGAGTCTTTTAGTGCC | ATGGATTCCTCTTTGTGCCATAG |
| GAPDH | TCGGAGTCAACGGATTTGGT | TTGGAGGGATCTCGCTCCT |

**Supplementary table 2. qRT-PCR primers.**
