## Supplemental table 3 for "Therapeutic Stress-induced Activation of PGCC Life Cycle Drives the Resistance Acquisition and Structured Tissue Differentiation"

| **Gene** | sequences (5'-3') |
| --- | --- |
| si-IL1α | GGUGUUCUCUGUUGCAGAAGUTT |
| si-IL1β | GCCAGGAUAUAACUGACUUTT |
| si-IL6 | GGAGACAUGUAACAAGAGUTT |
| si-IL8 | GCCAAGGAGUGCUAAAGAATT |
| si-CXCL1 | GCACAUCUGUUUUGUAACUTT |
| si-Control | UUCUCCGAACGUGUCACGUTT |

**Supplementary table 3. The sequences of small interfering RNA.**
